# Subcutaneous and orally self-administered high-dose carprofen in male and female mice: pharmacokinetics, tolerability and impact on cage-side pain indicators

**DOI:** 10.1101/2023.06.03.543582

**Authors:** Aylina Glasenapp, Jens P. Bankstahl, Heike Bähre, Silke Glage, Marion Bankstahl

## Abstract

Surgical interventions in mice are prerequisite in various research fields and require appropriate pain relief, not only to ensure animal welfare but also to avoid influence of pain on research findings. Carprofen is a non-steroidal anti-inflammatory drug that is commonly used as an analgesic for interventions inducing mild to moderate pain in animals. Despite its frequent use also in laboratory rodents, data on pharmacokinetics and side effects, and on its potential impact on behavioral pain indicators are rare.

This study aimed to determine pharmacokinetic and tolerability profiles of high dose carprofen in male and female C57Bl/6J mice, administered via single subcutaneous injection (s.c.) and oral self-administration per drinking water (d.w.). Plasma concentrations of carprofen were measured at various time points, and side effects were evaluated using a modified Irwin test protocol, hematology and histopathology. Additionally, potential effects on behavioral pain indicators commonly used to assess post-surgical pain, such as the mouse grimace scale, wheel running activity, burrowing, nesting and grooming behavior were investigated.

Quantification of carprofen in plasma revealed maximum plasma concentrations of 133.4 ± 11.3 µg/ml after 1 hour and an elimination half-life of 8.52 hour after single s.c. injection of 20 mg/kg carprofen. Oral self-administration of carprofen (25 mg/kg/24 h) resulted in a steady-state < 24 hours over 5 days after treatment start with plasma levels of around 60 µg/ml. The carprofen-medicated water was highly accepted, and increased d.w. intake was observed in the first 24 hours after exposure for both sexes (p < 0.0001). Irwin test detected only minor side effects, and hematology and histopathology where without pathological findings that could be attributed to carprofen treatment. Except for a decrease of 49-70 % in wheel running activity in male mice, behavioral pain indicators were only very mildly affected.

This study determined carprofen plasma levels in mice lying well above an estimated therapeutic concentration for both routes of administration. Carprofen was well tolerated at recommended high doses and may provide sufficient analgesia for minor interventions as well as be applied as a tolerable component in multimodal analgesic regimens.

## Introduction

Numerous experimental animal studies include surgical interventions, entailing the need for effective postoperative pain relieve. Nevertheless, structured literature reviews suggest that peri– and postoperative pain in laboratory rodents is often undertreated (1–3). Improvement of this situation is severely hampered by largely lacking pharmacokinetic (PK) and tolerability profiles in laboratory rodents even for standard analgesic drugs.

For analgesic treatment after surgical interventions in mice, an effective and well tolerable analgesic would be desirable, with stress-free administration for animals and convenient to apply in daily routine. Here, carprofen might be a promising analgesic candidate both for subcutaneous (s.c.) injection and for oral self-administration via drinking water (d.w.), as it is stable in water for at least 7 days (4) and there is no indication for aversion of rodents to carprofen medicated water (5, 6). Apart from its single usage it could also be a candidate for multimodal analgesia regimens, e.g. in combination with local anaesthesia and opioid drugs.

Carprofen is a nonsteroidal anti-inflammatory drug and widely used both for the management of chronic pain as well as acute soft tissue injury in animals. It is licensed for veterinary use as racemic mixtures of R– (–) and S– (+) enantiomers. Carprofen exerts its analgesic effects through inhibition of cyclooxygenase (COX) isomers, and species differences for its PK, COX-2 selectivity and enantiomer potency have been reported (7). So far, carprofen is often used to attenuate mild to moderate pain, but there is only limited knowledge about plasma concentrations and PK profiles of carprofen in mice, and so far only at doses that are considered to provide only limited analgesic efficacy in mice (4, 5, 8, 9). Data about tolerability and side effects are even more sparse, and there is a gap of knowledge concerning the analgesic efficacy of carprofen treatment in mice. Currently, only few reports on side effects are available for mice, but indicate that carprofen might be associated with an increased risk of gastrointestinal ulceration (10). After cardiac injury, increased inflammation reaction and disordered immune reaction under carprofen treatment are described (11, 12). Consequently, there is a need to investigate and describe side effects in a structured manner.

The use of behavioral indicators in mice to assess pain after surgical interventions has become increasingly important in research. The analysis of these parameters can improve not only animal welfare and minimize negative effects of pain on research outcome but may also provide reliable and objective measures of pain and analgesic efficacy. However, potential effects of carprofen on behavioral pain indicators, such as wheel running activity, burrowing, nesting and grooming behavior, or mouse grimace scale (13), have rarely been investigated in healthy mice so far (5). This knowledge is of particular importance in order to differentiate pain-induced behavior from drug side effects as a potential confounding factor.

The major aim of this study was to provide PK profiles and tolerability profiles for carprofen after s.c. and oral self-administration via d.w. in healthy mice at maximum recommended doses according to the current expert information on pain management for laboratory animals of the German Society of Laboratory Animal Science (GV-SOLAS; (14)). Moreover, the objective was to investigate the potential impact of carprofen treatment on established behavioral pain indicators that are commonly used to assess pain severity after surgical interventions.

## Material and methods

### General experimental design and blood sampling procedure

An overview of the experimental setup is provided in Fig 1. Animals were habituated to all handling procedures and interventions and were trained for burrowing before experiments started. Running disks were accessible constantly from beginning of habituation. During baseline phase 1 (BL1), data of all parameters of interest were obtained over 5 days, including food and water consumption, body weight, nest score, Irwin test, mouse grimace scale, body temperature, tail immersion test, wheel running and burrowing activity. This was followed by s.c. injection and subsequent blood collection for PK analysis at 1, 2, 3, 6, 12, 24, and 48 h post injection. After 2 weeks of recovery, a second baseline phase (BL2) was performed identical to BL1. Then, carprofen was administereds via d.w. for 5 days, and blood was collected at 3, 6, 12, 24, 36, 108, and 120 h after start of administration. Blood was sampled from a lateral tail vein by scalpel micro-incision, which has been shown to be of lower burden to the animals than withdrawal from the facial vein or the retro-orbital sinus (15). Blood was collected in EDTA tubes (Sarstedt 200 µL Microvette^®^, Germany) at all sampling time points. At each collection time point (see above), samples (∼ 120 µl) were taken from 6 mice (3 male, 3 female). Due to organizational reasons, number of animals is deviant for s.c. treatment after 1 h (2 male, 1 female), 2 h (4 male, 3 female), and 3 h (3 male, 5 female). No additional animals were added for 1 h, as an influence on outcome was considered irrelevant. Immediately after sampling, blood-containing tubes were placed on ice, then centrifuged (2500xG/10 min/4°C) and plasma was stored at –80°C until further processing. Directly prior to blood sampling, tests for tolerability (Irwin test) and analgesic efficacy (tail immersion test) were conducted in the respective 6 mice. At the end of the experiment, mice were killed by CO_2_ inhalation and subsequent cardiac puncture. Final blood samples were used for full blood count and analysis of electrolytes, glucose, and lactate. Further, autopsy of each animal including macroscopic investigation for abnormalities was performed. Stomach, small intestine, liver and kidneys were dissected for histological processing.

**Fig 1.**
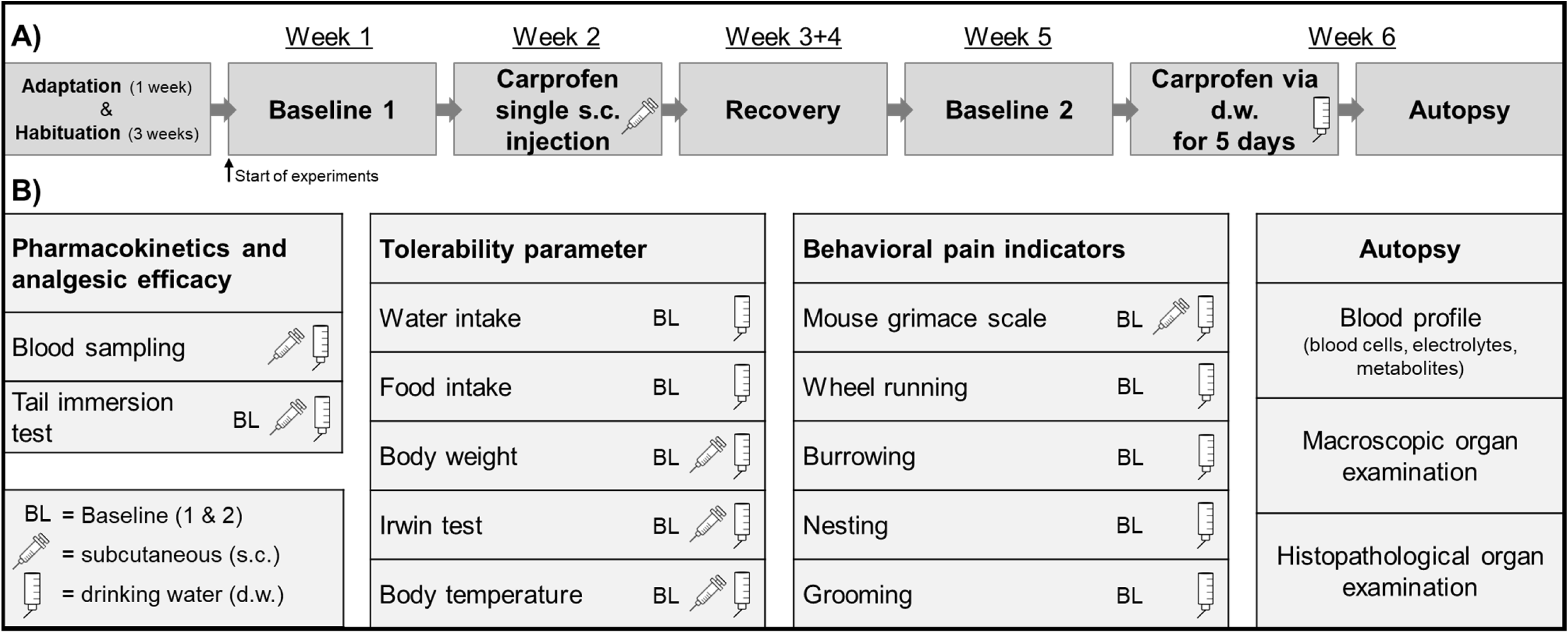
Experimental Design. A) Timeline of complete experimental setup. B) Overview of all interventions and parameters.

### Animals and cage set-up

This study was approved by the responsible state authority (Niedersächsisches Landesamt für Verbraucherschutz und Lebensmittelsicherheit under reference number 33.8-42502-04-21/3640). In-house bred C57Bl/6J mice were housed in groups of three individuals per cage (type 3, macrolone, UNO BV, The Netherlands) on standard bedding material (wood chips from spruce, poplar and aspen trunks, LAB.BED, Thomsen Räucherspäne Räucherholz GmbH & Co. KG, Germany) in a temperature-controlled facility under a 14 h/10 h light/dark cycle (lights on: 6:30 a.m.) with food (altromin 1320 standard diet, Altromin Spezialfutter, Germany) and filtered (particle filter, 5 µm pore size) tap water ad libitum. Experimental group size included 21 male and 21 female mice. As control for histologic analysis and blood profiles, 6 male and 6 female age-matched C57Bl/6J mice kept under the same conditions were used. At start of experiments with age of 12 weeks, mice weighed 28.0 ± 1.3 g (male) and 21.9 ± 1.2 g (female). Mice were transferred from the breeding area to the experimental room 4 weeks before start of the experiments. This time was subdivided into one week of adaptation phase to the new room without particular handling and three weeks of habituation to all handling procedures. Each mouse was adapted three times to weighing, manual fixation, restrainer fixation and tail immersion test (for tail immersion test habituation, water with body temperature was used). In their home cages, the mice had continuous access to nesting material, running disks installed on plastic houses, and wooden gnawing material as additional enrichment (plexx B.V. Aspen Bricks S, the Netherlands). Detailed cage and running wheel set-up is provided in S9 Fig.

### Analgesic

A commercially available carprofen solution (Zoetis, Rimadyl^®^ 50 mg/ml, injection solution for dogs and cats) was diluted with 0.9 % sterile sodium chloride (B. Braun, Germany) to achieve a 10 ml/kg injection volume. The s.c. injection dose was 20 mg/kg. For d.w. administration, the injection solution was diluted in filtered tap water targeting a carprofen dose of 25 mg/kg/24 h. Carprofen concentration calculation for group-housed mice (3 mice/cage, 7 cages per sex) was based on mean water consumption/24 h measured over 5 days (BL 2; male mice, 23.4 ml; female mice, 17.5 ml) and mean body weight (male mice, 29.9 g; female mice, 23.4 g). Carprofen-medicated d.w. was freshly prepared every morning in red polyphenylsulfon water bottles (Tecniplast, Italy). The carprofen dosage both for s.c. and oral administration was chosen based on the current GV-SOLAS expert information on pain management for laboratory animals (16).

### Sample preparation and liquid chromatography mass spectrometry analysis

Carprofen content in plasma was quantified by liquid chromatography mass spectrometry (LC-MS/MS). Plasma samples and analgesic calibrators (25 µL) were thawed and diluted using 100 µL extraction solvent (acetonitrile/ methanol 1/2) containing 12.5 nM efavirenz as internal standard (equals a final concentration of 10nM/sample) in a 1.5 mL reaction tube (SafeSeal®, Sarstedt, Germany) for analyte extraction and protein denaturation. Samples were mixed for 30 sec using a vortex mixer and frozen overnight at −20 °C to complete protein precipitation. Then, samples were thawed and centrifuged for 10 min at 20,800 x g/4 °C for protein separation. For mass spectrometry analysis, samples were diluted 1:500 using dilution solvent (acetonitrile/methanol/water 2/2/1) containing 10 nM internal standard, and 100 µL were transferred into mass spectrometry vials (Wicom, Heppenheim, Germany) with inserts (Macherey-Nagel, Düren, Germany) for carprofen quantification. This involved chromatographic separation on a reversed phase C18-column (ZORBAX Eclipse XDB-C18 1.8 µ, 50 x 4.6 mm, Agilent, Santa Clara, California, USA) connected to a C18 security guard (Phenomenex, Aschaffenburg, Germany) which was kept on 40°C during the whole analysis. A linear gradient was performed using an HPLC-system consisting of two LC-30AD HPLC pumps, a SIL-30AC temperature controlled autosampler, a DGU-20A5 degasser, a CTO-20AC oven, and a CBM-20A control unit (Shimadzu, Duisburg, Germany). 10 µL of the sample was injected. Elution started with 80% of solvent A (water + 0.1 % formic acid). Within 7 minutes, the amount of solvent B (methanol 0.1% formic acid) was linearly increased to 95%. This composition was maintained for 3 minutes followed by a 3 minute re-equilibration of the column back to 80/20 (solvent A/solvent B). The total analysis time was 13 minutes at a flow rate of 0.4 mL/min. The retention time of carprofen as well as efavirenz (internal standard) was 7.7 min. Analytes were detected by triple quad mass spectrometry (5500QTRAP; Sciex, Framingham, Massachusetts) in multiple reaction monitoring (MRM) mode. Ionization was achieved using negative electrospray ionization at 650°C. For carprofen, the mass transition m/z 272 → 226 was optimized for quantification, and for efavirenz, the mass transition m/z 314 → 244 was used. Control of HPLC and the mass spectrometer as well as data sampling was performed by Analyst software (version 1.7., Sciex). For quantification, calibration curves were created by plotting peak area ratios of carprofen, and the internal standard versus the nominal concentration of seven calibrators containing 15.6 nM – 1000 nM carprofen prepared in mouse plasma. The calibration curve was calculated using quadratic regression and 1/x weighing.

Pharmacokinetic analysis was performed by using Phoenix WinNonlin (Version 8.2, Certara, Princeton, NJ). Noncompartmental analysis was applied for determination of plasma elimination half-life of carprofen.

### Tolerability assessment and side effects

#### Irwin test and mouse grimace scale

Prior to blood sampling, tolerability of carprofen was examined by a modified Irwin test (17) at 1, 2, and 3 h after s.c. injection and at 3, 6, 12, 24, 36, 108, and 120 h after start of voluntary uptake via d.w.. The test included for observations without handling, in the open field, and with handling (S1 Table). Parameter assessment was subdivided into several categories, such as excitation, coordination, sedation, and autonomous system. Mouse grimace scale was scored as an additional parameter of the Irwin test assessment, and visually assessed (without handling of the mice) in the home cage (18, 19) according to established mouse grimace scale score (20). Immediately after Irwin test procedure, rectal body temperature was measured (PhysioSuite^®^ for mice and rats, Kent Scientific Corporation, USA). For visualization of Irwin test, a heat map was created showing percentage of changes per investigated time point after treatment (n= 3 male and 3 female per time point, except s.c. treatment after 1 h (2 male, 1 female), 2 h (4 male, 3 female), and 3 h (3 male, 5 female). For further analysis of the Irwin test outcome, a sum score of each individual mouse and test category (excitation, coordination and sedation) was generated. Total score values were added for a total sum score. Negative score values were handled as positive score values for addition to sum score. Test parameters for which presence or absence were recorded were given 2 points if they deviated from normal. Handling associated vocalization was compared to animal individual BL and assessed with 2 points when deviant from BL. Vocalization during reflex testing (eyelid, pinna) was scored with 2 points when present. Presence of unusual behavior in free text (fasciculation) was scored with 2 points per observed behavior.

#### Food and water intake, and clinical score

Food and water consumption were gravimetrically measured every 24 h per cage during baseline phases (BL1 & BL2). During substance administration via d.w., water bottles were additionallyweighed at blood sampling time points (for respective cages), resulting in fluid consumption data for day and night. Clinical score was determined twice a week during baseline (BL1 & BL2) phases and daily during carprofen administration phases. Clinical scoring included judgement of activity, general state of health, behavior, body posture and body weight (S1A Fig).

### Pain indicators other then grimace scale

A variety of behavioral parameters, such as burrowing, nesting, and grooming behavior, and wheel running activity have been suggested as potential indicators of pain in laboratory mice (13). They were included in this study to determine potential impact of analgesic treatment on their outcome. Their usefulness as pain indicators may depend on the specific type and intensity of pain, and further research is needed to fully evaluate their reliability and validity. However, several studies demonstrated practicability and sensitivity of these parameters after surgical interventions to provide an overall perspective on general health status (5, 21, 22). To avoid impact of cage change on behavioral pain indicators, cages were changed already 2-3 days prior to start of assessment (BL1 & BL2, carprofen via d.w.). Nesting and gnawing material were provided continuously throughout the whole experiment.

#### Wheel running

Commercially available angled mouse running discs and igloos were purchased from ZOONLAB GmbH (Castrop-Rauxel, Germany). Running time (h:min:sec), running distance (km), average velocity (km/h) and maximum velocity (km/h) were recorded as previously described (22–24) by bike computers (Sigma BC 16.16 STS, SIGMA-ELEKTRO GmbH, Germany). In brief, the sensor was fixed on the outside of the cage in close proximity (<1 cm) to the magnet (neodymium, 10 x 3 mm), which was glued to the running disc. To avoid any accidental shifting of the igloos by the mice, the position of the setup in the cage was secured by a customized pedestal (polyethylene) which was attached to the cage floor with adhesive tape (S9 Fig.). Wheel running parameters were continuously recorded and read out in the morning between 8-9 am and in the afternoon between 4-6 pm. Mice had continuous access to running discs during the complete experimental period. A wide range of habituation and training effects, and variability during experiments of running activity is described (22). The first BL measurement in this study was performed 1 week after equipment of home cages with running disks. Data were recorded twice for baseline (BL1 & BL2) and under carprofen treatment. Only the second baseline (BL2) recording was used as reference for data analysis.

#### Burrowing

Burrowing procedure was performed to investigate motivated goal-directed behavior (13) during substance administration according to a modified protocol from Deacon et al (25). For assessment of burrowing behavior, empty transparent bottles (volume 330 ml, length 15 cm, diameter 5 cm, polycarbonate) were filled with food pellets (altromin 1320 standard diet, Altromin Spezialfutter, Lage, Germany). First, mice were habituated twice to burrowing overnight by placing food-pellet filled bottles in the front left corner of the cage. Next morning, bottle and food were removed. For experiments, bottles were filled with 151 ± 2 g (mean ± SD) food pellets, placed in the cage and removed after 2 h. Latency to start of burrowing (min) was visually measured for the first 30 min. In cases where the mice started burrowing later, 30 min were used as value for data analysis. At the end of burrowing procedure (after 2 h) the remaining pellets in the tube were weighed and the amount burrowed (%) calculated. Burrowing procedure was started between 11 am and 1 pm. Two baseline data sets (BL1 & BL2) were recorded for each cage, mean of both was used for statistical analysis and is labeled as BL. During oral substance administration, burrowing behavior was assessed every day for 2 h (at 24 h, 48 h, 72 h, and 96 h).

#### Nest building

Nesting behavior was observed to detect alterations of intrinsic motivated behavior under drug treatment according to established nesting score (5): 1 = no cotton pieces grouped together; 2 = cotton pieces paired together in one or two pairs; 3 = 3 cotton pieces grouped together; 4 = all cotton pieces grouped together; 5 = all 4 cotton pieces grouped together and completely shredded. Assessment of nest building test was performed using 4 cotton rolls (Ø 12 mm x 37 mm, ANT Tierhaltungsbedarf, Germany) per cage (S10A Fig). Existing nesting material was taken out and 4 new cotton rolls were provided 10 h after start of carprofen treatment via d.w. and nest score was recorded for the first time 2 h later. Subsequently, nests were scored every day in the morning (8-9 am; 14, 38, 62, 86, 110 h after administration of nesting material) and in the afternoon (4-6 pm; 24, 48, 72, 96 h after administration of nesting material). Two individual baselines (BL1 & BL2) of nesting were performed, for statistical analysis median and range of both is displayed and labeled as BL.

#### Grooming transfer test

Grooming behavior as potential indicator for pain or distress was evaluated according to Oliver et al. (5). Two baseline (BL1 & BL2) trials were performed. For statistical analysis, median and range of both is displayed and labeled as BL. At 10 h after start of carprofen administration via d.w., 8 µL of fluorescent GloGerm Oil (Glo Germ Company Utah, USA) were applied to the neck region of the mice (onto the skin). Using an UV flashlight (UVL 1006, Glo Germ Company Utah, USA), 2 h after application and then everyday in the morning between 8-9 am (14, 38, 62 h after application of fluorescence) fluorescence distribution was scored according to an established grooming score (5) (S10B Fig): 1 = Fluorescence is strong at the application site on the forehead between the ears; 2 = Fluorescence is present at the application site and front, and/or rear nails; 3 = Fluorescence is at the application site and the ears, signal may be present at front and/or rear nails; 4 = Fluorescence is absent from the nails and ears but traceable amount remains at application site; 5 = Fluorescence is no longer visible. Two female mice with alopecia in the neck region were excluded from analysis.

### Analgesic efficacy

To evaluate antinociceptive efficacy of carprofen, two baselines (BL1 & BL2) of hot water tail immersion test were performed for each individual mouse. After carprofen treatment, the test was performed prior to all blood sampling time points (3 male and 3 female mice per time point). A water bath maintained constant temperature at 50.1 ± 0.3 °C (n = 167) for heat stimulus. During the procedure, mice were placed into a costumized cylindric restrainer (polyvinyl chloride) equipped with a cut-out for the mouse tail. The distal 3 cm of the tail was marked and subsequently immersed in hot water and time manually stopped (stop watch ROTILABO^®^, Carl ROTH, Germany; seconds, two decimal places) until reflex reaction. This was repeated three times with less than 20 sec interval between measurements (26). The mean of the three measures was used for statistical analysis. To prevent potential heat-induced skin damage when showing no reflex reaction, the tail tip was removed from hot water after 10 sec cut-off time. Difference of latencies between treatment and individual baseline were normalized and displayed as ratio to individual baseline. Comparing baseline measurements between sexes, a distinctly longer latency time to tail withdrawal of 2.8 ± 0.7 s for male mice versus 1.7 ± 0.5 s for female mice (BL1, p < 0.0001) was measured (data not shown).

### Final blood analysis

After CO_2_ inhalation, blood from final cardiac puncture was sampled in EDTA tubes (Sarstedt 500 µL Microvette^®,^ Germany) and immediately stored on ice until analysis, to generate full blood count (scil Vet abc, scil animal care company GmbH, Germany) including leukogram of white blood cells (WBC), lymphocytes (LYM), monocytes (MO) and granulocytes (GRA). Red blood cell (RBC) count includes hemoglobin (HGB), hematocrit (HCT), platelets (PLT), mean corpuscular volume (MCV), mean corpuscular hemoglobin (MCH), mean corpuscular hemoglobin concentration (MCH), red cell distribution width (RDW), and mean platelet volume (MPV). In parallel, additional blood analysis provided status of electrolytes (concentration of Na^+^, Ca^2+^, Cl^-^), metabolites (glucose and lactate concentration) and blood (hemoglobin, hematocrit) (ABL815 Flex blood gas analyzer, Radiometer, Denmark). Here, blood after cardiac puncture was sampled in capillaries (safeCLINITUBES, Radiometer, Denmark) and stored at room temperature (< 2 h) until analysis.

### Autopsy and histology

Directly after cardiac puncture, a routine dissection and visual inspection of the organs in the thoracic and abdominal cavity was performed. Thereafter, stomach, duodenum, jejunum, liver and kidneys were immediately removed and fixed in formaldehyde solution (4 %). The small intestine was flushed with sodium chloride (0.9% Braun, Germany) and prepared in an improved swiss-roll technique (27). After at least 3 days of formaldehyde fixation, tissues were trimmed according to the RITA-Guidelines (28–30), dehydrated (Shandon Hypercenter, XP), and subsequently embedded in paraffin (Histoplast, Thermo Scientific). Sections (2 – 3 μm thick, microtome Reichert-Jung 2030) were deparaffinized in xylene and H&E stained according to standard protocols. From each cage, organs of one individual mouse were randomly chosen for histopathological analysis (n = 7 male, 7 female). A trained veterinary pathologist performed blinded evaluation (Axioskop 40, Carl Zeiss AG, Germany) and scoring of the sections (stomach, intestine). The kidneys and liver were evaluated for overall pathological findings. The scoring (31) of the small intestine and stomach considered four, respectively three general criteria: (1) inflammatory cells (2) intestinal architecture, (3) degree of ulceration, if present, and (4) percentage of area involved.

The criterion (1) inflammatory cells was subdivided into evaluation of severity and the maximal extent of the inflammatory cells regarding the histologic layers of the mucosa and graded from 0 – 4 for each with a maximum score of 8. For scoring of the intestine, the mucosal architecture was likewise subdivided in evaluation of the epithelial and the mucosal layer and graded from 0 – 4 for each with a maximum score of 8. The degree of ulceration included score values from 0 – 3 and the area involved score values from 0 – 4. All criteria per organ were added up, resulting in an overall maximum score of 23 for the intestine and 15 for the stomach. Images of histological sections were taken on an Axioskop 40 microscope utilizing the software ZEN (Carl Zeiss AG).

### Statistical analysis

Animal-individual data were recorded for carprofen plasma levels, body weight, clinical score, grooming activity, Irwin test parameters, mouse grimace scale, body temperature, tail immersion test, hematology, and histology, whereas data for food and water consumption, nesting, burrowing, and wheel running activity were recorded on cage level. Results are presented as mean and data points (plasma concentration-time curves); mean ± standard deviation (carprofen intake, wheel running, burrowing, tail withdrawal latency, body temperature, body weight, blood parameters, fluid and food consumption); median and individual data points (sum score Irwin test); or median and range (nest score, grooming score). Comparisons of two groups were performed by two-tailed Student’s t-test (blood parameters) or Wilcoxon test (body temperature). Multiple comparisons were analyzed by one-way ANOVA (burrowing, tail withdrawal latency, grooming) or Friedman’s test (nesting) and Dunnett’s or Dunn’s multiple comparisons test, respectively, or two-way ANOVA and Šídák’s multiple comparisons (wheel running). GraphPad Prism 9 (Version 9.3.1, GraphPad Software, USA) was used for all statistical analyses. Unless stated otherwise, a p-value < 0.05 was considered statistically significant.

## Results

### Favorable half-life of carprofen after s.c. injection and stable plasma concentrations during oral intake via d.w

To generate PK profiles of carprofen in mice, tail vein blood was collected at 1, 2, 3, 6, 12, 24, and 48 h after s.c. injection and at 3, 6, 12, 24, 36, 108, and 120 h after start of oral self–administration. Single s.c. administration of 20 mg/kg carprofen resulted in maximum plasma concentrations of 133.4 ± 11.3 µg/ml after 1 h and an elimination half-life of 8.52 h (Fig 2). Plasma concentrations were above an assumed therapeutically useful level of 24.3 µg/ml, representing the in vitro canine whole blood assay IC_80_ value for COX-2 inhibition (7), for up to 24 h after injection. Oral self-administration of carprofen (25 mg/kg/24 h) started in the morning of the light phase and resulted in a steady-state < 24 h after treatment start with a plasma level of around 60 µg/ml (Fig 3A). Maximum plasma concentrations of 93.0 ± 30.6 µg/ml were observed after 24 h. Carprofen-medicated water was well accepted and resulted in increased drinking water intake compared to BL, especially in the first 24 h after start of treatment for both sexes (p < 0.0001) (Fig 3B). During the first 24 h, male mice consumed 79% and female mice 86% more water compared to BL2 measurements (S5A Fig). The increased water intake resulted in an ingested carprofen dose of 43.6 ± 7.2 mg/kg (males 44.8 ± 7.1 mg/kg, females 42.4 ± 7.7 mg/kg) within the first 24 h, and intake of the target dose (25 mg/kg/day) was almost reached at 12 h after treatment start (S2A Fig). From 72 h to the end of the experiment, water intake was stable for both sexes and resulted in dose intake of approximately 30 mg/kg/24 h (Fig 3B). Regarding circadian rhythm, mice consumed between 49 % (0-24 h) and 35 % (96 – 120 h) of total carprofen intake during the light phase (S2B Fig). No obvious differences between female and male mice were noted for plasma levels resulting from both routes of administration.

**Fig 2.**
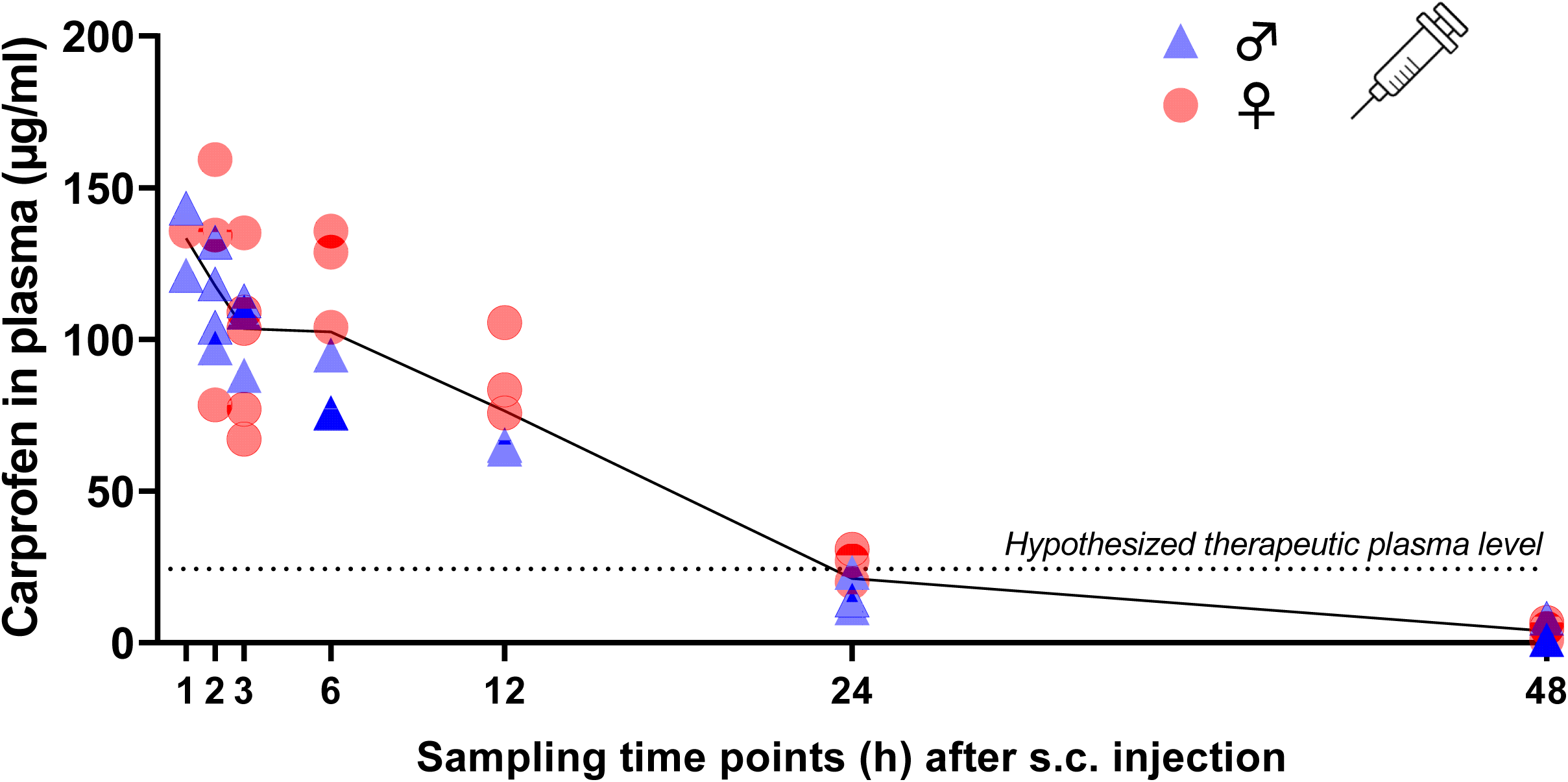
Plasma concentration-time curve following single s.c. injection of 20 mg/kg carprofen. Individual data points (n = 6, 3 male and 3 female per time point; exceptions: s.c. 1 h n = 2 male, 1 female; s.c. 2 h n= 4 male, 3 female; s.c. 3 h n = 3 male, 5 female) and mean of all animals per time point (black line) are presented. Assumed minimal therapeutic plasma level (dotted line) is displayed according to (7).

**Fig 3.**
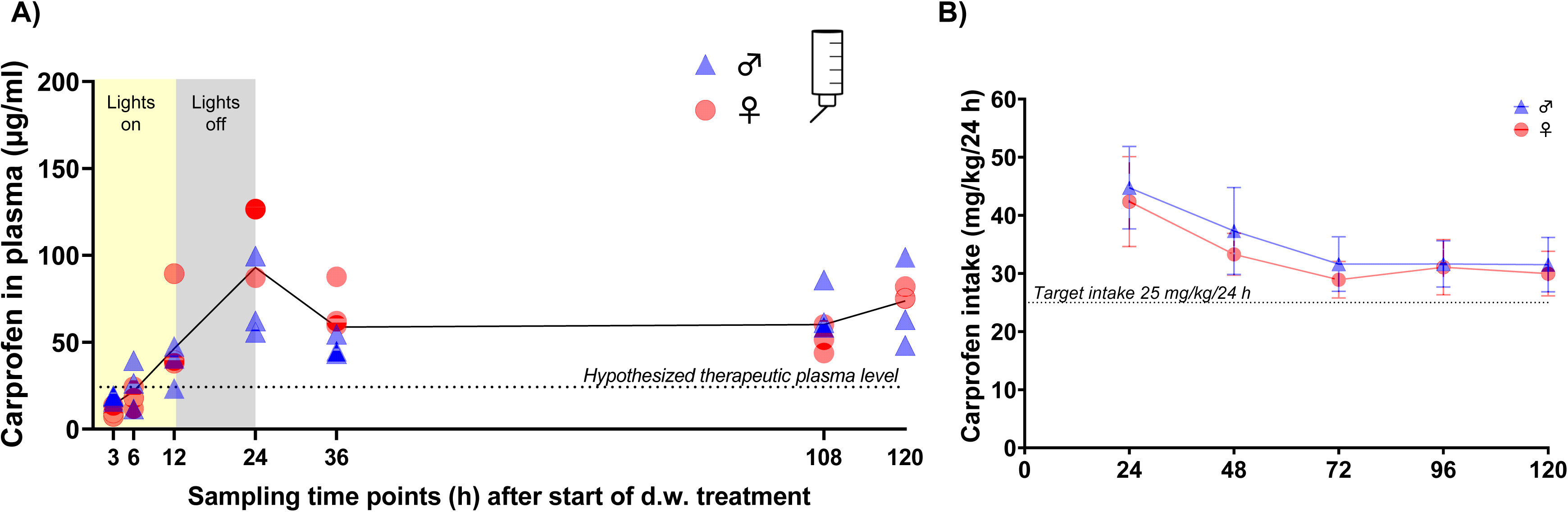
Carprofen plasma levels and dose intake during oral self-administration. A) Plasma concentration-time curve during oral intake of carprofen (intended dose: 25 mg/kg/24 h) via drinking water (d.w.) over 5 consecutive days. Individual data points (n = 6, 3 male and 3 female per time point) and mean of all animals per time point (black line) are presented. Assumed minimal therapeutic plasma level (dotted line) is displayed according to (7). B) Carprofen intake (mg/kg) is displayed for 24 h intervals after start of oral treatment and is above target intake dose of 25 mg/kg/24 h (dotted line). Dose intake was calculated by consumption and concentration of carprofen-medicated drinking water.

### Carprofen is well tolerated after s.c. injection and d. w. administration

Tolerability and potential side effects were evaluated by a modified Irwin test protocol including a variety of read out parameter (S1 Table). Investigated parameters were categorized in excitation, coordination, sedation, autonomic symptoms, and mouse grimace scale and displayed in a heat map (Fig 4A). After s.c. injection, both sexes showed a slight decrease in grip strength, whereas male mice also showed slightly reduced locomotion/exploration. During oral intake, in both sexes excitation was observed, represented by increased vocalization during reflex testing (eyelid, pinna) and by increased tail elevation (straub-like). However, handling-induced vocalization was reduced. Coordination, visual placement, tail suspension test and grip strength were reduced in some individuals. Pointing to a sedative effect, decreased locomotor activity was observed. For the categories excitation, coordination, and sedation, a sum score for each individual animal and time point is presented in Fig 4B. After s.c. injection, increased score values for coordination and sedation were observed, whereas during d.w. treatment significant changes compared to baseline in all three categories were present. However, deviations in score values were mainly expressed in a low grade of Irwin score (S1 Table).

**Fig 4.**
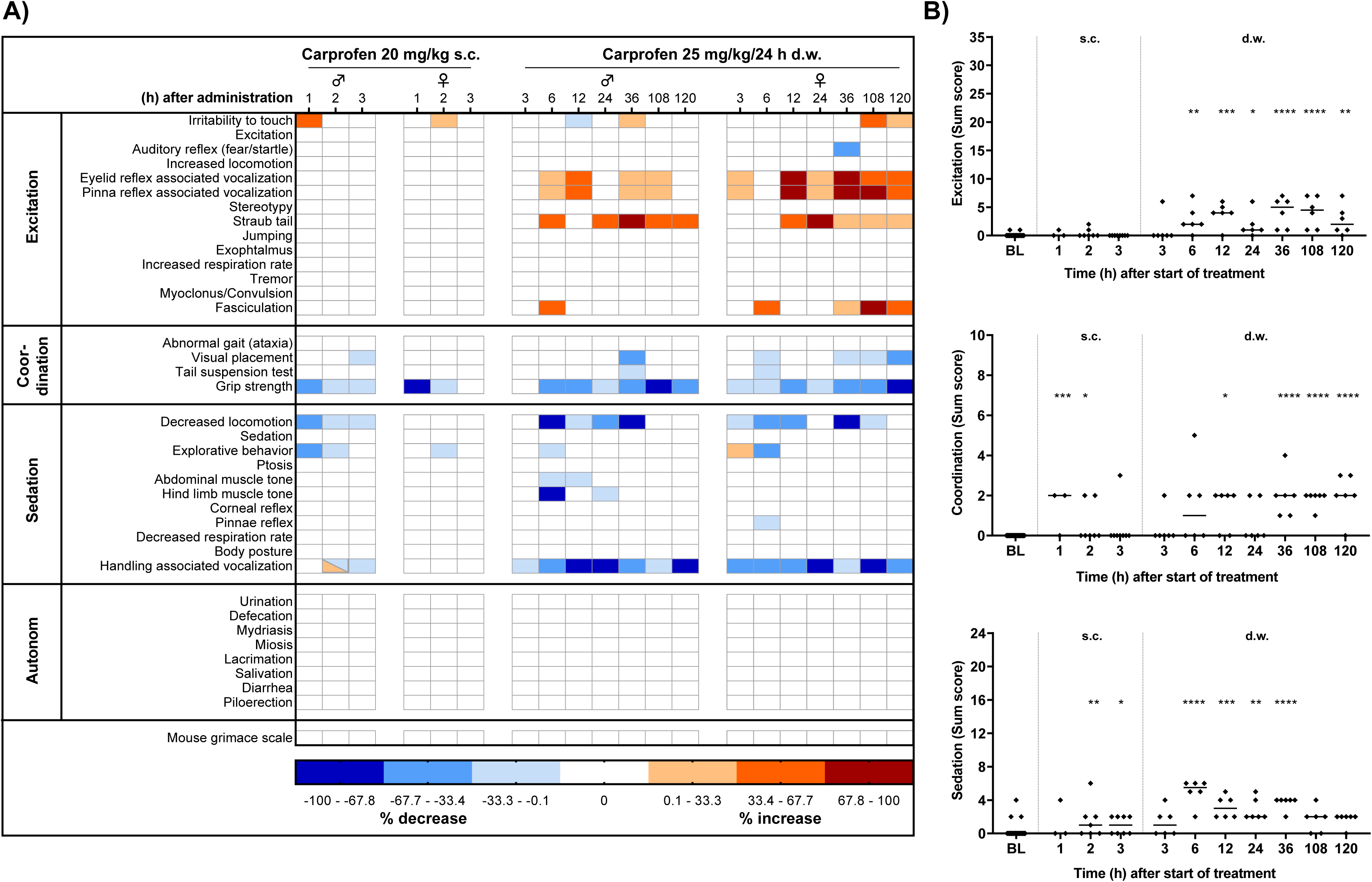
Tolerability of carprofen assessed by modified Irwin test. A) Side effects of carprofen on behavioral parameter are displayed as heat map. Per time point and sex, 3 mice were investigated (exceptions: s.c. 1 h n = 2 male, 1 female; s.c. 2 h n= 4 male, 3 female; s.c. 3 h n = 3 male, 5 female). Increase or decrease is presented as % change of total number of animals per sex and timepoint. B) Kruskal-Wallis test followed by Dunn’s multiple comparisons test show significantly higher sum score values compared to baseline in the categories excitation, coordination and sedation (*p < 0.05; **p < 0.01; ***p < 0.001; ****p < 0.0001).

Live scoring of the mouse grimace scale (9) was performed as part of Irwin test procedure (Fig 4A) in the home cage and did not reveal increased score values at any time point.

For rectal body temperature, no sex differences were observed comparing individual baselines (BL1 male vs. female, p = 0.1619; BL2 male vs. female p = 0.1676). However, a significant difference for female mice between first and second baseline measurement was present (p = 0.0002; S8B Fig). Therefore, respective BL1 and BL2 data were used for statistical analysis of carprofen’s potential impact on body temperature (Fig. 5). After s.c. carprofen injection, no differences in body temperature compared to individual baseline were monitored (p = 0.4763) (Fig 5A). However, significantly increased body temperature was observed during d.w. treatment (p < 0.0001) (Fig 5B).

**Fig 5.**
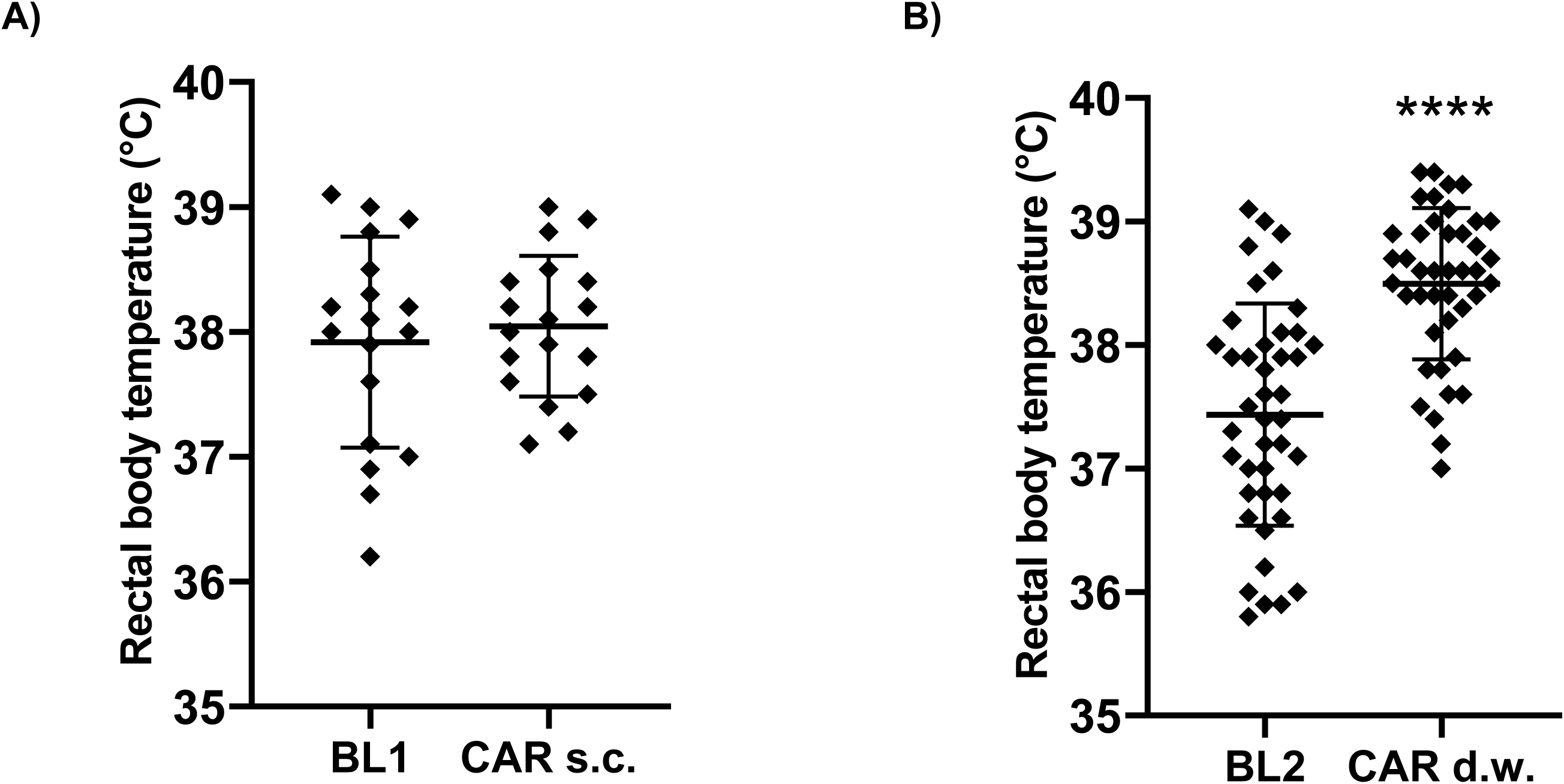
Carprofen intake via d.w. influences body temperature. Data from all measurements after s.c. carprofen treatment (A) and intake via d.w. (B) are shown. Wilcoxon test compared paired data after s.c. (n = 18; 9 male, 9 female) and d.w. (n = 42; 21 male, 21 female) treatment and shows increased body temperature during d.w. treatment, but not after s.c. injection of carprofen compared to individual baseline (****p < 0.0001).

A clinical score (S1A Fig) was assessed twice per week during baseline phases and daily during s.c. (until day 2 after injection) and d.w. treatment including activity, general condition, behavior, body posture and body weight. In only 2 female and 1 male mice, minor and transient increases in clinical score were observed during carprofen treatment (S1B Fig).

The body weight during experiments was stable in both sexes (S4 Fig), but female mice exhibited a slight decrease in food consumption 120 h after start of oral carprofen administration via d.w. (p = 0.0086) (S5B Fig).

### Impact on behavioral pain indicators: Oral carprofen administration has none to minor impact on burrowing, nesting, and grooming behavior or mouse grimace scale, but sex-dependently affects wheel running activity

Wheel running activity was measured over five consecutive days during dark and light phases to detect potential influence of analgesic treatment on voluntary exercise. Male mice showed a significant decrease both in running time overnight (min) after 48 h (–70 %; p < 0.0001), 72 h (–55 %; p = 0.0002), and 120 h (–49 %; p = 0.0048) and running distance overnight (km) after 48 h (–75 %; p = 0.0001), 72 h (–55 %; p = 0.0021), and 120 h (–52 %; p = 0.0315) during oral analgesic administration vs. respective time points during BL2 (Fig 6A). Female mice displayed comparable wheel running activity to BL2, except for a decrease in running time overnight (min) after 48 h (–25 %; p = 0.0168) (Fig 6B). For both sexes, running time, distance and velocity within the light phase were not affected by analgesic treatment.

**Fig 6.**
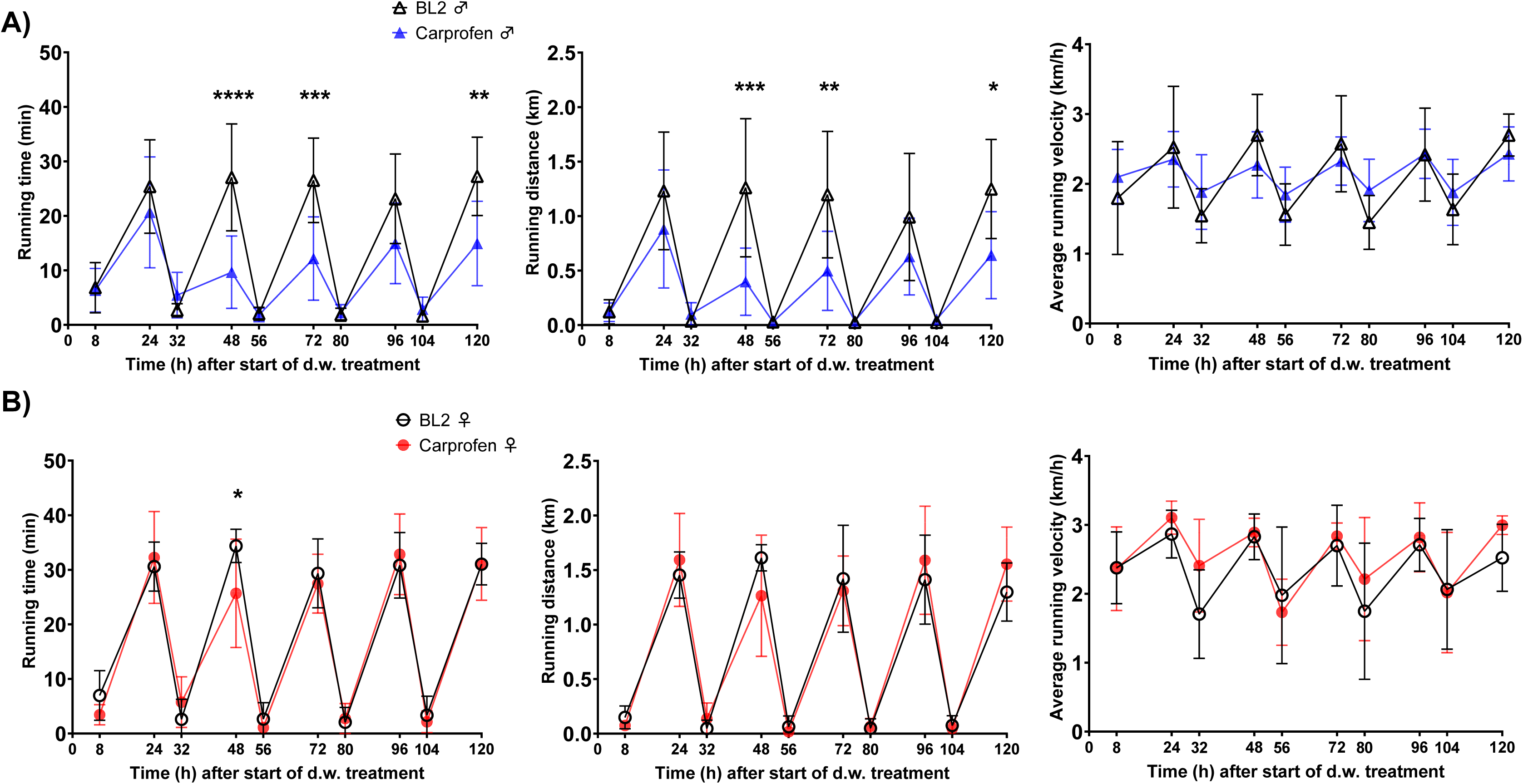
Carprofen treatment via d.w. is associated with decreased wheel running activity in male mice. A) Running time (min), running distance (km) and average running velocity (km/h) is shown normalized to total time (h) during read out intervals (8 h during day and 16 h during night) for A) male and B) female mice. Running wheel data were documented between 8 and 9 am in the morning and between 4 and 6 pm in the afternoon. Data are shown as mean ± SD (n = 7 cages per sex, 3 mice/cage). Two-way ANOVA and Šídák’s multiple comparisons test reveals decrease in wheel running time and distance in male mice under carprofen treatment (*p < 0.05; **p < 0.01; ***p < 0.001; **** p < 0.0001).

During carprofen administration via d.w., burrowing performance was assessed every day (in 24 h intervals) after start of treatment. During the burrowing time of 2 h, male mice burrowed continuously stable amounts of pellets both at baseline assessments and after analgesic administration (Fig 7A). Burrowing latency was significantly shorter after 96 h under carprofen medication (p = 0.0453) (Fig 7B). For female mice, we observed distinctly higher fluctuation in burrowing activity (latency and amount burrowed) both during baseline phases and after substance administration (Fig 7A,B).

**Fig 7.**
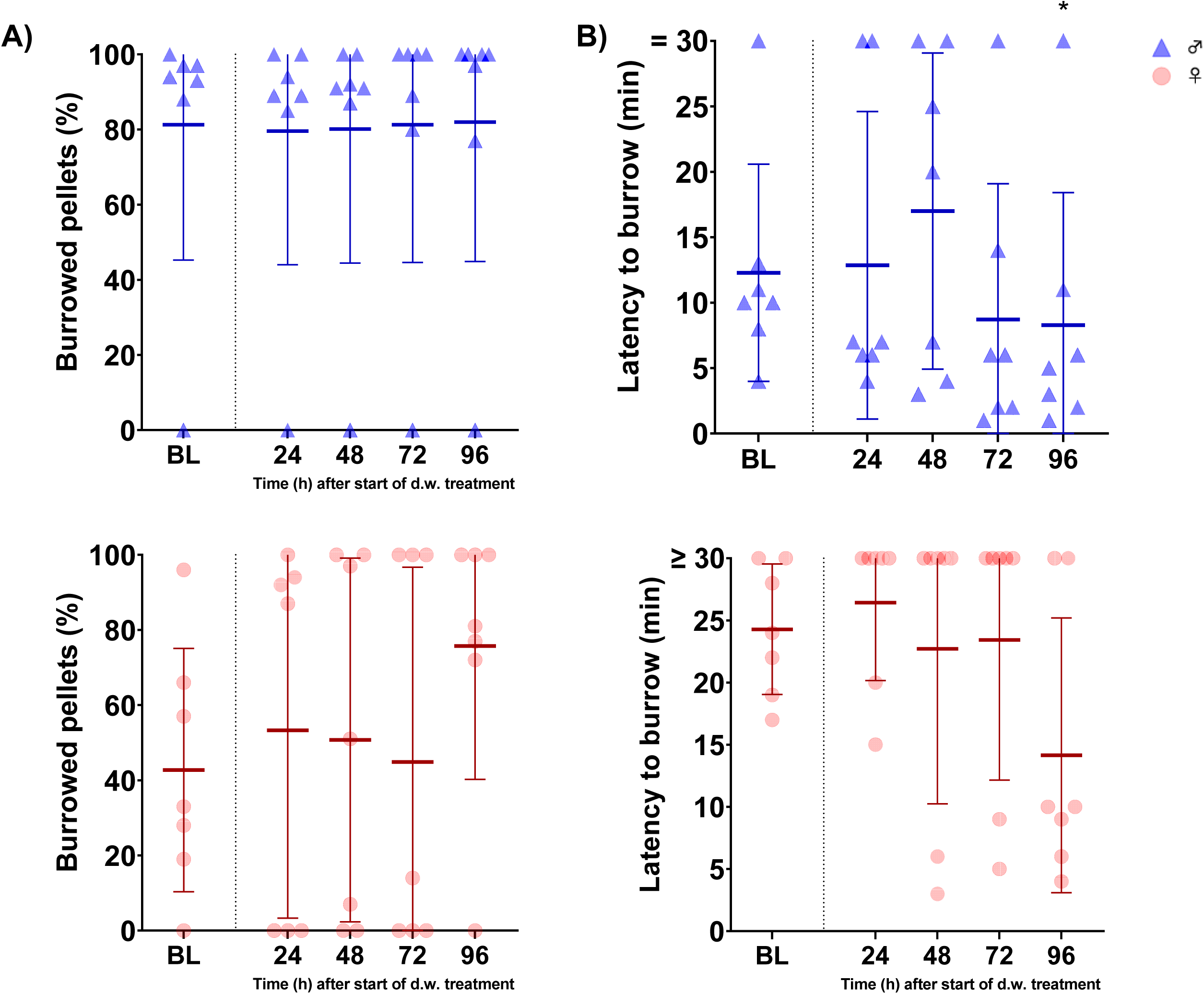
Burrowing behavior is not influenced by carprofen treatment via d.w.. A) Latency time to start of burrowing behavior after placing the burrowing tube in the cage for male (top) and female (bottom) mice, n = 7 cages per sex, 3 mice/cage. Data are shown as mean ± SD. One-way ANOVA and Dunnett’s multiple comparisons test reveal shorter latency to burrow in male mice 96 h after treatment vs baseline (*p < 0.05). B) No changes in volume of burrowed food pellets (in % after 2 h for male (top) and female (bottom) mice, n = 7 cages per sex, 3 mice/cage) were observed.

Nesting behavior was assessed over 5 consecutive days twice per day using an established nest score (5). Nest scores under carprofen treatment developed comparable to median of BL 1 and 2 in mice of both sexes (Fig 8A). Time to achieve nest score 5 was unchanged for males, but shorter for females under carprofen medication (Fig 8B).

**Fig 8.**
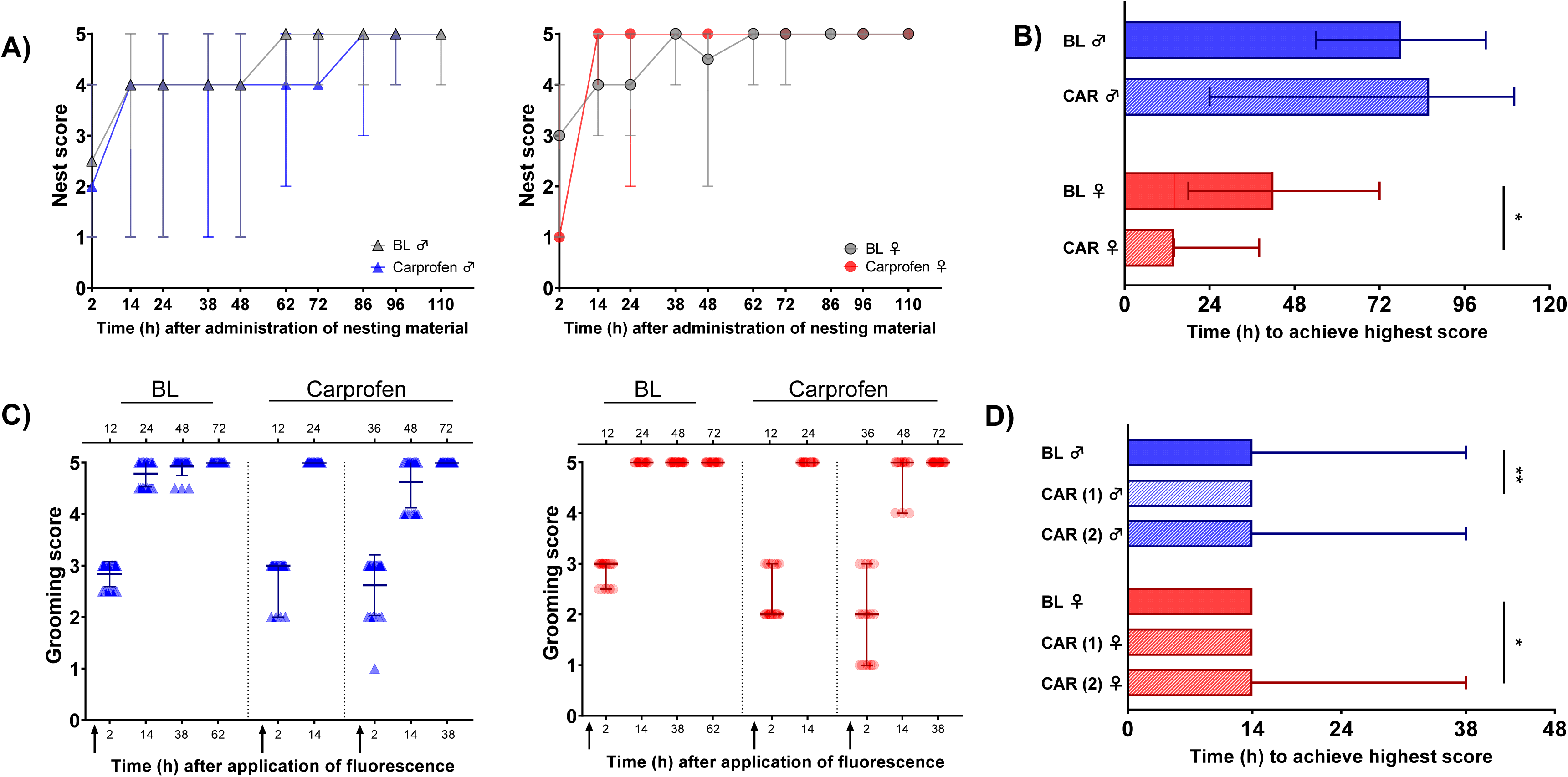
Nesting behavior remains unaffected and grooming activity slightly prolonged during carprofen treatment via d.w.. A) Nest scores after administration of nesting material for male (left) and female (right) mice. Nesting material was provided 10 h after start of d.w. treatment. Data are presented as median and range (n = 7 cages per sex, 3 mice/cage). Friedman’s test und Dunn’s multiple comparisons test did not reveal significant differences at any time point. B) Time (h) after administration of nesting material until nest score 5 was achieved is displayed. Data are presented as median and range (n = 7 cages per sex, 3 mice/cage). Wilcoxon test demonstrates significantly shorter time to score 5 (p = 0.0313) for female mice during d.w. treatment. C) Grooming scores of individual animals are shown for male (left) and female (right) mice as median and range (n = 21 per sex). Fluorescence suspension was applied 10 h after start of d.w. treatment to mouse skin. Upper x-axis indicates time points during BL and after start of d.w. treatment (Carprofen). Lower x-axis shows time points after administration of fluorescence suspension. Arrows indicate administration of fluorescence. D) Time (h) after application of fluorescence to achieve grooming score 5. Data are presented as median and range (n = 21 mice per sex). One-way ANOVA and Dunnett’s multiple comparisons test shows significantly shorter time to achieve score 5 for male mice under carprofen d.w. treatment vs. BL in the first trial (p =0.0042). Kruskal-Wallis test and Dunn’s multiple comparisons tests reveals prolonged time to achieve score 5 for female mice in the second trial (p = 0.0488).

Grooming activity was assessed under an UV light at 2 h, 12 h, 36 h and 60 h after start of carprofen d.w. treatment according to established scoring (5). Grooming test started by application of fluorescent oil to the neck region between the ears at 10 h after oral treatment start. Under carprofen treatment, the oil was applied twice. Grooming activity, represented by score values, developed similar under treatment compared to median of BL 1 and 2 (Fig 8C). However, a slightly shorter time to achieve score 5 was observed for male mice in the first trial under carprofen treatment (p = 0.0042) (Fig 8D) whereas female mice showed slightly prolonged grooming behavior in the second trial (p = 0.0046).

### No anti-nociceptive effect of carprofen in the hot water tail immersion test

Hot water tail immersion test was performed to test antinociceptive efficacy by latency to tail withdrawal from hot water (Fig 9). Calculation of withdrawal latency as ratio to baseline did not reveal statistically significant anti-nociceptive effects in the chosen test paradigm after s.c. (Fig 9A) and d.w. administration (Fig 9B). A correlation analysis of carprofen concentration in plasma and tail withdrawal latencies was performed but did not show statistical correlation (S3 Fig).

**Fig 9.**
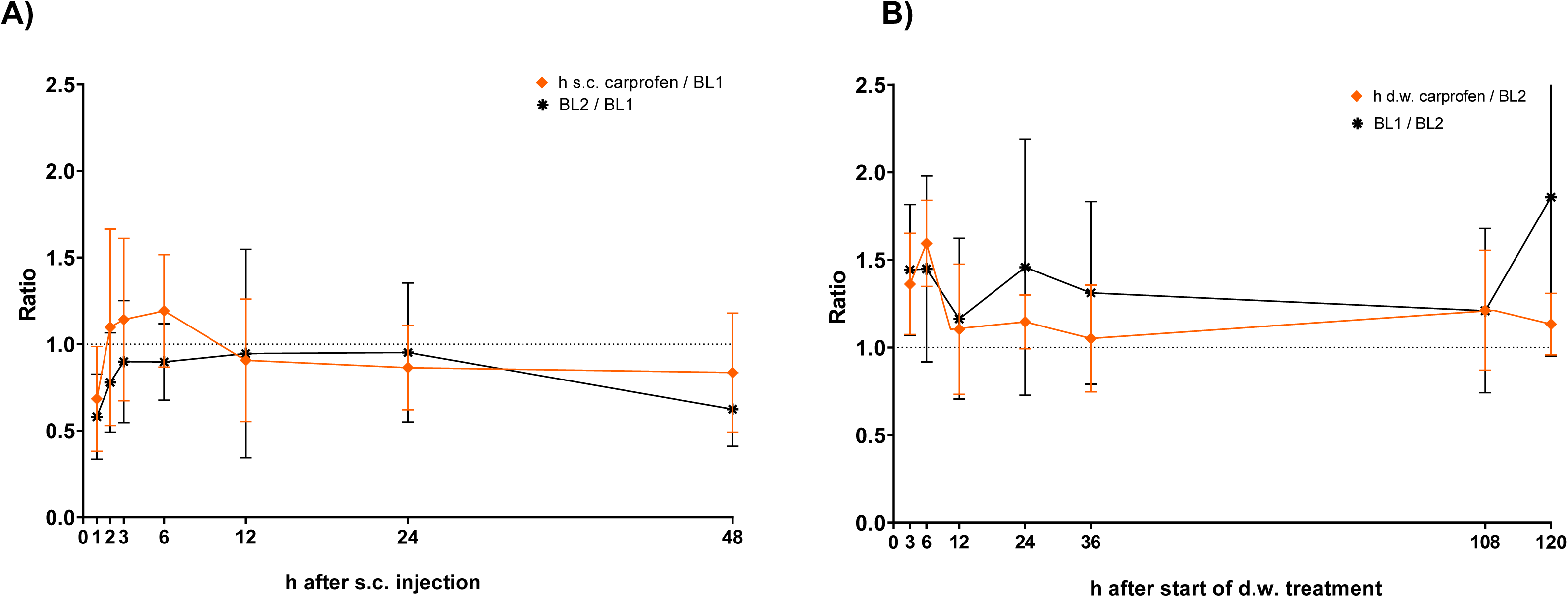
Analgesic efficacy of carprofen after s.c. injection and during treatment via d.w.. Latency to tail withdrawal in the hot water tail immersion test at respective time points after A) single s.c. injection and after B) oral d.w. treatment. Latency to tail withdrawal is displayed as ratio to animal individual BL. For comparison, the ratio of BL latency data is displayed. Latencies were calculated as mean of total three measures per animal and time point. Per time point, n = 6 mice (3 male, 3 female) are shown (exceptions: s.c. 1 h n = 2 male, 1 female; s.c. 2 h n= 4 male, 3 female; s.c. 3 h n = 3 male, 5 female).

### Comprehensive blood analysis and histological examination does not indicate adverse effects of carprofen

To assess potential side effects of continuous carprofen treatment for 5 days via d.w., organs of interest (stomach, duodenum, proximal part of jejunum, liver, and kidneys) were examined in detail by a veterinary pathologist. Representative images of histologic stomach and jejunum sections from control and carprofen-treated mice are shown in Fig 10. No tissue damages or signs of inflammation were observed, and rated histology score values (control: 0.30 ± 0.87 (range 0-3) n = 12; carprofen 0.14 ± 0.53 (range 0-2) n = 14; data not shown) of stomach and small intestine were comparable between groups.

**Fig 10.**
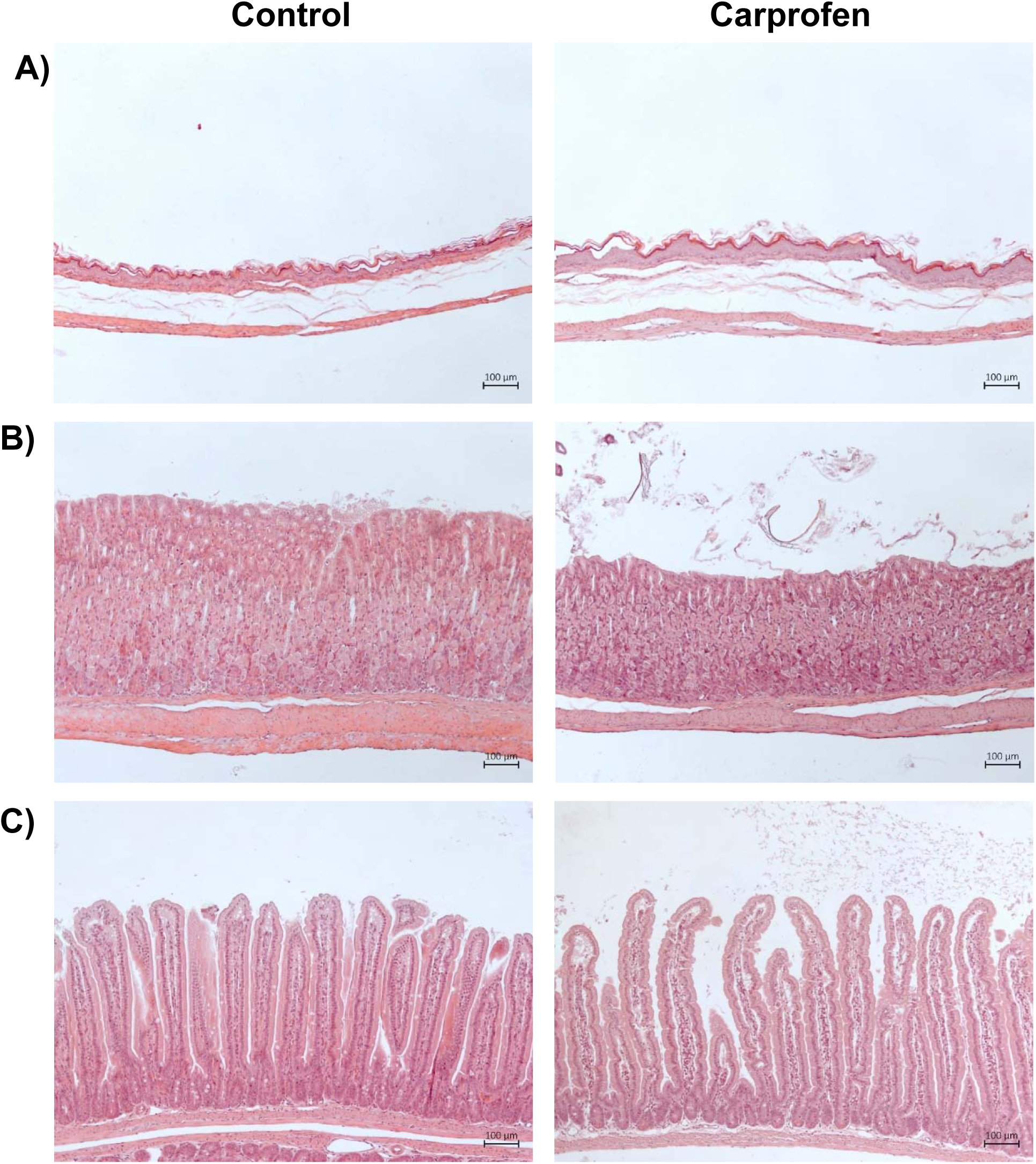
Carprofen has no adverse effects on histology of selected organs. Representative hematoxylin-eosin staining of A) forestomach B) glandular stomach and C) proximal part of jejunum of control animals (left) and after 5 days of carprofen d.w. treatment (right).

Standard blood profile after cardiac puncture was measured after 120 h of continuous carprofen d.w. treatment and compared to healthy control animals. Leukogram of white blood cells (WBC), lymphocytes (LYM), monocytes (MO), and granulocytes (GRA) as well as red blood cell (RBC) count including hemoglobin (HGB), hematocrit (HCT), platelets (PLT), mean corpuscular volume (MCV), mean corpuscular hemoglobin (MCH), mean corpuscular hemoglobin concentration (MCH), red cell distribution width (RDW), and mean platelet volume (MPV) is provided in S6 Fig. Unpaired t test revealed lower LYM cell count (10^3^/mm^3^) (control vs carprofen, p = 0.0156), HGB (g/dl) (control vs carprofen, p = 0.0019), HCT (%) (control vs carprofen, p = 0.0022), and MCH (pg) (control vs carprofen, p = 0.0003). Electrolyte and metabolite data did not indicate deviations between control and carprofen-treated animals (S7 Fig). However, significantly lower HGB and HCT in carprofen-treated mice were confirmed by measurement in a second system.

## Discussion

### Study objective and main findings

As basis for evidence-based development of post-operative pain management using carprofen, this study aimed to determine its pharmacokinetics, tolerability, and impact on pain indicators in C57Bl/6J mice after s.c. injection and oral self-administration via d.w.. Following a single s.c. injection of 20 mg/kg carprofen, C_max_ was reached before 1 h, and an elimination half-life (t_1/2_) of 8.52 h was determined. Oral self-administration of 25 mg/kg/24 h resulted in a steady-state plasma concentration within 24 h, which was maintained during the complete experimental phase of 5 days. Carprofen-medicated water was well accepted with even an increased fluid intake during analgesic administration over 5 days, especially in the first 48 hours. Moreover, carprofen was well-tolerated, with only minor and transient side effects observed for both routes of administration. The behavioral pain indicators were not influenced (grimace scale) or only very mildly, in the form of slightly accelerated burrowing and nest-building or slightly accelerated (male mice) or delayed (female mice) grooming behaviour. However, carprofen administration for 5 days via d.w. resulted in decreased wheel running activity, seen mainly in male mice, of up to 70 %. Hematological and histopathological examination did not provide evidence for any pathological effects on organs.

### Carprofen, current dose recommendations, and PK data

Carprofen is a non-steroidal anti-inflammatory drug (NSAID) that is commonly used in laboratory rodents to relieve pain and inflammation. It is approved for clinical use in veterinary patients in the EU (various species) and the US (dog) in multiple preparations, but application for pain relieve in rodents is off label. For mice, dose recommendations of carprofen range between 2-20 mg/kg s.c. with an injection interval of 12-24 h (10, 32). According to the current expert information “pain management for laboratory animals” of the German Society for Laboratory Animal Science (GV SOLAS), 5-20 mg/kg carprofen s.c. with an injection interval of 12 h is recommended for pain relief in mice (14). In terms of the length of time carprofen remains above the assumed minimum plasma concentration and its rather slow elimination a 12 h injection interval appears reasonable. However, analgesic efficacy after s.c. injection of 5 mg/kg is controversially discussed. Studies using single injection of 5 mg/kg indicate that there might not be sufficient analgesic coverage after surgery (33, 34), whereas repeated injections might alleviate mild post-surgical pain (35). Recently published results indicate adequate pain relief of s.c. injected carprofen in doses of 20-25 mg/kg as single analgesic after surgery assessed by mouse grimace scale (9, 36).

As current literature research indicates, evidence-based dose recommendations for oral self–administration of carprofen in mice are also rare. GV SOLAS currently suggests doses of 10–25 mg/kg for oral administration via d.w. (16). While assessment by mouse grimace scale has indicated an analgesic effect of 25 mg/kg/24 h via d.w. after craniotomy (36), 30 mg/kg/24 h had only minimal benefit after laparotomy in mice (5). Nevertheless, the non-invasive and stress-free nature of medication via drinking water makes this approach favorable. Moreover, carprofen is reported to be stable in drinking water without adverse effects on consumption (4). Having these heterogeneous findings and recommendations in mind, our study examined the maximum recommended dose from the GV SOLAS for both administration routes.

Reported PK data showed that 5 mg/kg carprofen s.c. resulted in a maximum plasma concentration (C_max_) of 52.5 µg/ml and t_1/2_ of 6.88 h in female CD1 mice (8). Oral intake of 10 mg/kg/24 h via d.w. resulted in C_max_ of 32 µg/ml after 24 h in female CD1 mice (5) and in 17.0 ± 2.9 µg/ml after 12 h in male C57Bl/6J mice (4). In our study, we determined a t_1/2_ of 8.52 h after s.c. injection (20 mg/kg) with plasma levels above an assumed therapeutic threshold (see below) for up to 24 h, and relatively stable plasma concentrations of around 60 µg/ml from < 24 h up over 5 days during oral intake over d.w.. However, in vivo therapeutic plasma levels of carprofen in mice are still unknown to date. Current literature often refers to an estimated therapeutic threshold above 24 µg/ml calculated from in vitro data using canine blood cells (corresponding to a carprofen concentration inhibiting 80 % of COX-2 activity) (7). Although carprofen plasma levels were clearly above this estimated threshold, we could not find a correlation between plasma levels and latency of tail withdrawal in the present study. Here it is important to mention that not only the plasma level might be considered as reference for therapeutic efficacy, but also analgesic concentration in the tissue and receptor binding/enzyme inhibition, both of which were not determined. Further, the applied tail immersion test, although having been shown to respond to NSAIDs (37–39), is much more sensitive for detection of anti-nociceptive effects by opioids.

### Tolerability profile of carprofen

Overall, only minor side effects were observed by the Irwin test, hematology, and histopathology, suggesting a good tolerability of carprofen in the investigated dose in mice. Evaluation of potential side effects of analgesics is important as they might put extra burden to the animal but also impact outcome measures of the scientific question. For tolerability assessment, close monitoring of the animal after drug administration is necessary. Therefore, in this study, apart from clinical scoring and body weight measurement, a modified Irwin test protocol (17) including 38 parameters in total was performed to assess potential carprofen-induced excitation/sedation, or impact on coordination or autonomous function. Additionally, hematologic and histopathologic evaluation was conducted in order to obtain a comprehensive impression of drug impact when continuously administered for 5 days, a period of time that may be required for the management of moderate to severe post-operative pain. In the Irwin test, minor impact on coordination and slight excitatory and sedative effects were observed during prolonged treatment via d.w.. Main changes were increase of reflex-associated vocalization, decrease of drip strength, decreased locomotion, and decreased handling-associated vocalization. In general, however, mice were in a very good overall condition and side effects were so mild that they did not become obvious during clinical scoring. Behavioral changes are rarely reported in veterinary patients (40).

Studies about carprofen effects on hematologic parameters in mice are sparse. In our study, blood count after 5 days of oral treatment revealed reduced lymphocyte concentration and tendency towards reduced white blood cells compared with control mice. Interestingly, a study demonstrated that carprofen (5 mg/kg s.c.) increased blood leukocyte numbers (monocytes) and leukocyte migration in male C57BL/6 mice in a model of venous thrombosis (41), while other hematological parameters were unaffected. We also found decreased HGB and HCT values, which are most likely a consequence of blood sampling during our experiments (42).

However, some NSAIDs are associated with anemia due to gastrointestinal bleeding (43) or induced hemolytic anemia (44) in humans. Additionally, carprofen at dose of 20 mg/kg is potentially associated with a risk of gastric ulceration in mice (10). In our experiment, the detailed histologic examination of a representative part of the experimental group (n = 14/42 mice) did not suggest any adverse effect of carprofen on stomach, duodenum, proximal part of jejunum, liver or kidneys after chronic d.w. treatment over 5 consecutive days when compared to untreated control mice. In line with our findings, a recently published study investigated possible toxic effects of 20 mg/kg carprofen on female CD1 mice by analyzing serum chemistry, fecal occult blood, and histopathological evidence of gastritis. Their findings showed that carprofen was not associated with toxic renal or hepatic effects and suggest tolerability of repeated s.c. injections every 24 h up to 7 d (45).

### Impact of carprofen on behavioral pain indicators

Analgesics might inadvertedly influence the outcome of pain indicators commonly used after surgical interventions, such as burrowing, nesting and grooming behavior, or wheel running activity (13), which would limit their validity during post-operative pain assessment. On the other hand, awareness of this influence would allow considering this in the assessment.

Burrowing and nest building are described as non-invasive and sensitive indicators of pain, distress and suffering in mice, as these parameters can reflect their well-being and motivation (21). Mice with pain or under stress tend to burrow less and show a decrease in nesting performance compared to healthy mice (46). It was shown that burrowing in carprofen-treated mice (5 mg/kg) started earlier after laparotomy compared to controls without pain medication (47), supporting its utility for detecting analgesic efficacy as well. Here, we detected a slightly accelerated start of burrowing behavior on the fifth day of analgesic treatment in male mice, which could be due to carprofen. Instead, it might suggest that despite extensive exposure to burrowing material, habituation, had not yet been fully completed, meaning that even more extensive familiarization would be necessary to achieve stable measurement values for this pain indicator.

It is reported that a change in nesting behavior can detect pain up to 24 h after laparotomy by scoring of nesting consolidation (5). The same study established grooming activity as a pain indicator and revealed sensitivity of grooming score up to 48 h following surgery. During carprofen administration via d.w., we found a slightly reduced time to reach the highest nest score in female mice compared to BL conditions. Other than for burrowing behavior, here, we can exclude habituation effects as the mice had continuous access to nesting material since they were born. Whereas nesting is to monitor without disturbance of the animals, grooming activity (as assessed in our study) requires additional restraining and handling of the animal to administer the fluorescent oily suspension and to score the grooming activity. In our study, we applied the fluorescent suspension twice during carprofen treatment and found slightly decreased (male mice) or slightly increased (female mice) times to achieve the highest grooming score in one out of two trials, respectively.

Voluntary wheel running activity has been suggested as a potential measure of pain in mice, because it is a natural behavior that is shown voluntarily when mice are provided access to a running wheel and can be easily monitored, objectively measured and quantified (22). It was shown to be sensitive to detect distress in a mouse model of colitis (48), inflammatory (49) or neuropathic pain (50). After surgical removal of the mammary fat pad, wheel running activity was reported to be reduced at 24 h independent of saline or analgesic treatment with carprofen, buprenorphine, or both (33). The authors concluded that the used opioid on the one hand had a sedative effect associated with approximately 80% less running activity whereas the applied carprofen dose of 5 mg/kg (s.c.) had no sufficient analgesic effect being associated with a running activity decrease of approximately 60%. Our finding of decreased running time and distance (mainly in male) mice under carprofen treatment via d.w. underlines that knowledge about analgesic influence on outcome measures is highly relevant for data interpretation. Reduced wheel running activity after surgery in carprofen-treated mice must not necessarily mean insufficient pain relieve but might just be explained by carprofen effects on this pain indicator. However, it is important to note that wheel running behavior may be sensitive to further factors other than pain or analgesic drugs, such as cage change, presence of experimenters in room, handling of the animal or blood sampling. For example, we observed increased running activity in the first afternoon both during baseline phases and treatment, which might be associated with presence of experimenter in the room or a consequence of more frequent handling of the animals at the respective days.

### Analgesic efficacy of carprofen in mice

Evidence-based dose regimens for carprofen to successfully reduce pain as well as plasma levels for therapeutic efficacy in mice are not known yet. The hot water tail immersion test is a commonly used method to assess thermal nociception and was included in our study to inform us about anti-nociceptive efficacy of carprofen. The test is based on the principle that exposure to a noxious thermal stimulus, such as hot water, will elicit a reflexive withdrawal response, which will be prolonged by analgesics impacting thermal nociception (38, 51). The type of restrainment during the test (38) and the number and interval of repetitions (26) is known to influence the outcome of the test. Also a sex difference in thermal nociception in mice has been reported (52), which was confirmed by our results, as males exhibited longer latencies during baseline measurements compared to females. Besides, the outcome can depend on water temperature. For example, the test successfully detected antinociceptive effects of the NSAIDs diclofenac, etocolac, and indomethacin in mice at a water temperature of about 52 °C (53, 54), whereas ibuprofen treatment prolonged tail withdrawal latency only at 45 °C but not at 50 °C (51). As ibuprofen and carprofen are related substances, we might have found a more pronounced antinociceptive effect at a lower water temperature. Moreover, potential influence of stress-induced analgesia by handling or restraining should be taken into consideration (55). To avoid this confounding effect, mice were habituated extensively to all handling procedures including bathing the tail before start of experiments. So far, according to current literature research, our study is the first to investigate the analgesic efficacy of carprofen using the hot water tail immersion test. However, despite comparatively high carprofen doses applied, no antinociceptive effect was found for both routes of administration, compared to individual baseline measurements. For future studies evaluating NSAIDs we will therefore include additional tests potentially better reflecting also the analgesic effects based on anti-inflammatory action.

Nevertheless, there is evidence that carprofen doses of 20-25 mg/kg s.c. effectively reduce postoperative pain after laparatomy when the mouse grimace scale is used as pain indicator (9). After craniotomy, doses of 10 and 25 mg/kg both s.c. and oral per d.w. demonstrated a significant reduction in mouse grimace scale scores, where injected carprofen was more effective immediately after surgery, and oral carprofen treatment was more effective in females via self-administration (36). After laparotomy, also doses of 20 and 25 mg/kg as s.c. single injection are reported as effective post-operative pain relief utilizing the mouse grimace scale (9). In view of the carprofen plasma concentrations determined here, this suggests that the hypothesized therapeutic threshold of 24 µg/ml is not set too high. Effective pain relief with a dose of 5 mg/kg is discussed, as some studies reported effective analgesia after surgical intervention (35), whereas the majority of studies did not observe clear pain relief (33, 34, 47, 56). However, exact dose and dosing regimen required to provide sufficient pain relief can also vary depending on multiple factors such as the mouse strain, sex, age, the experimental procedure, and the individual animal’s health status. Moreover, carprofen’s analgesic effect is mainly mediated through its anti-inflammatory mechanism of action and will therefore not address all levels of pain perception, suggesting that a combination with analgesics of a different mechanism of action is a more promising approach for strengthening pain relief than further dose increase.

### Limitations and future directions

Limitations of the present study must be considered when interpreting the results. Firstly, it is important to note that measurement of the plasma concentration of carprofen may not necessarily reflect the concentration at the target receptor, and thus might not always be indicative for therapeutic efficacy. Furthermore, despite inclusion of a test for acute anti-nociception, the levels of plasma concentration required for therapeutic efficacy in mice after surgery remain to be determined in a future study. It is also important to mention that behavioral parameters can be influenced by a variety of factors other than analgesic treatment. In the context of our study particularly the blood sampling as most invasive procedure should be noted. Inclusion of a control group undergoing blood sampling but not analgesic treatment might therefore also have added information for interpreting the impact on pain indicators, especially wheel running and grooming activity. However, the study demonstrates that even high doses of carprofen do not induce relevant adverse effects, suggesting that a respective dose adaptation may be considered by experimenters and veterinarians for future post-surgical application of carprofen as mono-analgesic. Moreover, continuous administration of carprofen via d.w. can be considered a promising refinement approach for the post-operative phase, provided that d.w. consumption is measured. As d.w. consumption is often reduced in the early post-operative phase, a combination of administration routes (starting s.c., continuing via d.w.) also seems a practical approach. Future investigations will focus on the use of carprofen in the context of surgical interventions, as well as exploring alternative pain models that better reflect clinical conditions. Furthermore, carprofen may be of high interest as part of a multimodal approach for pain management in mice.

## Conclusion

PK and tolerability profiles of high doses of carprofen in mice support suitability of both s.c. and oral self-administration in the early post-operative phase. While assumed therapeutic plasma concentrations were achieved, only minor side effects were detected, no pathological findings were observed in hematology or histopathology, and apart from wheel running activity, behavioral pain indicators remained largely unaffected. The results of this study demonstrate that carprofen is well tolerated at high doses in mice and is therefore a promising candidate to be used in a multimodal analgesic approach.

## Acknowledgements

The authors thank the animal keepers and laboratory members of the Institute for Laboratory Animal Science and Central Animal Facility for their support. We are especially grateful to Viktor Lutscher from the pathology working group and Annette Garbe from the Metabolomics Core Facility for excellent technical assistance.

---

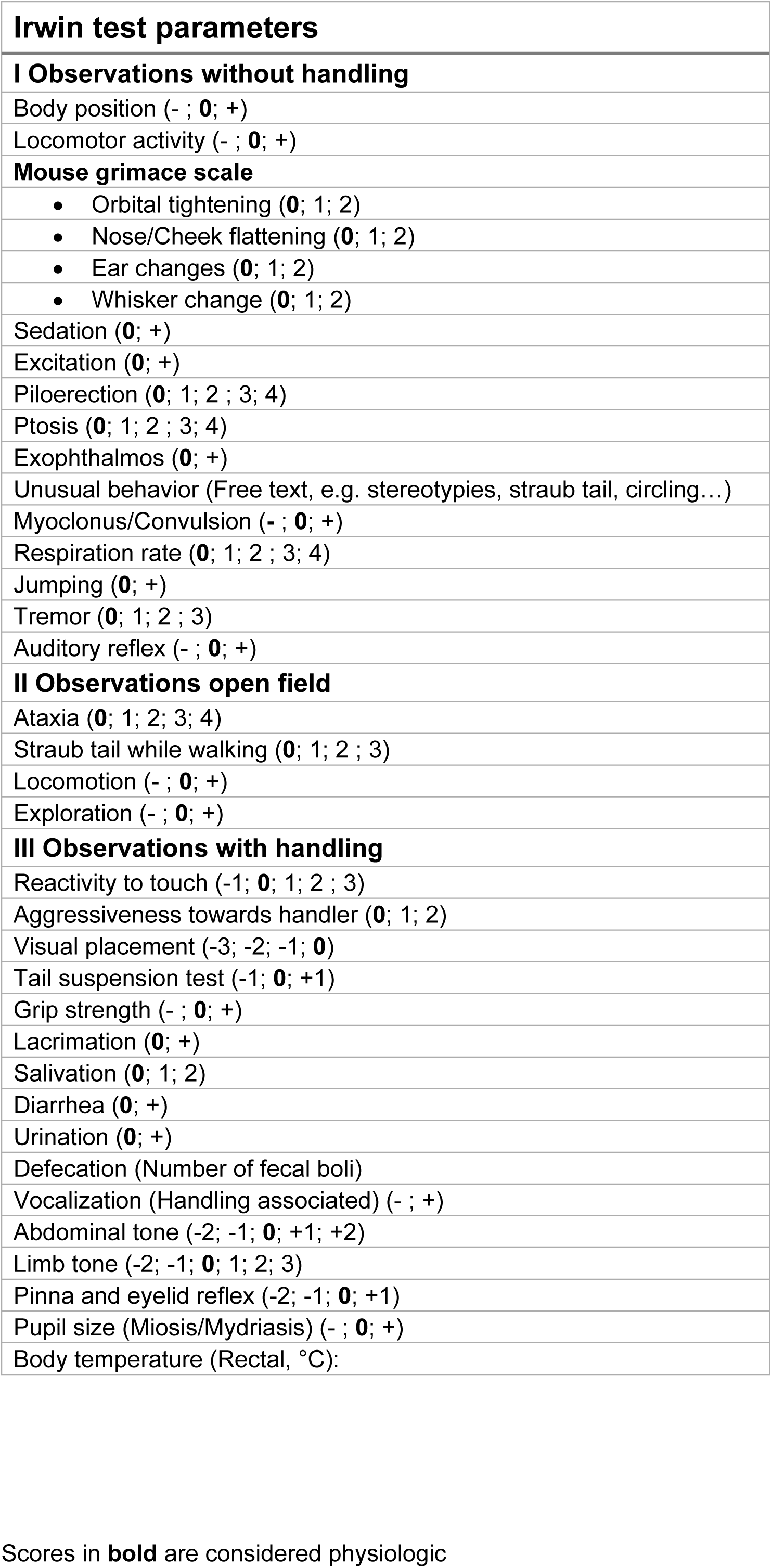

