## Supplemental data for "Subcutaneous and orally self-administered high-dose carprofen in male and female mice: pharmacokinetics, tolerability and impact on cage-side pain indicators"

**S1 Fig. Clinical score.** A) Clinical score was assessed daily during s.c. and d.w. treatment. B) After s.c. injection, 1 male mouse showed an increased clinical score value after 24 and 48 h. During d.w. treatment, 2 female mice expressed an increased clinical score value at 48 h.

**S2 Fig. Carprofen intake during light and dark phase.** A) Carprofen intake (mg/kg) within the first 24 h during d.w. treatment. Dose was calculated from individual water intake and concentration per cage and sex. B) Carprofen intake (%) during the light and dark phase is displayed for 12 h intervals of total intake within 24 h (mean).

**S3 Fig. Carprofen plasma levels are not associated with latencies to tail withdrawal.** Latency difference of tail immersion test (%) shows no correlation (Spearman correlation) to level of carprofen in plasma (nM). Simple linear regression (black line) describes the relationship between both variables graphically.

**S4 Fig. Development of body weight.** Daily body weight of A) male and B) female mice throughout the experimental course. Data points represent individual mice (n = 21 per sex), mean ± SD is shown. During baseline phases (BL1, BL2), mice were weighed 2x/week.

**S5 Fig. Fluid and food consumption. C**onsumption (g) of drinking water (d.w.) and carprofen-medicated water (d.w. + carprofen) is presented over 5 consecutive days, respectively. B) Food consumption (g) during baseline and carprofen treatment is presented over 5 consecutive days. Data are shown as mean ± SD (n = 7 cages/sex; n = 3 mice/cage).

**S6 Fig. Blood count.** Full blood count of naïve control mice (n = 6 male, 6 female) and carprofen treated mice (n = 21 male, 21 female) was measured after final cardiac puncture. Unpaired t test was used to test for differences (*p < 0.05, **p < 0.01, ***p < 0.001, **** p < 0.0001). Dotted lines represent reference values provided by manufacturer (scil vet, Vet abc). A) Leukogram of white blood cells (WBC), lymphocytes (LYM), monocytes (MO), and granulocytes (GRA). B) Red blood cell (RBC) count includes hemoglobin (HGB), hematocrit

(HCT), platelets (PLT), mean corpuscular volume (MCV), mean corpuscular hemoglobin (MCH), mean corpuscular hemoglobin concentration (MCH), red cell distribution width (RDW), and mean platelet volume (MPV).

**S7 Fig. Blood analysis of electrolytes and metabolites.** Blood analysis of naïve control mice (n = 6 male, 6 female) and carprofen-treated mice (n = 21 male, 21 female) was performed after final cardiac puncture. Unpaired t test was used to test for differences (*p < 0.05, **p < 0.01, ***p < 0.001, **** p < 0.0001). Concentrations of electrolytes, glucose, lactate, hemoglobin (HGB) and hematocrit (HCT) are shown.

**S8 Fig. Body temperature.** Time course of rectal body temperature (°C) during baseline phases, as well as after s.c. and during d.w. treatment shown for A) male and B) female mice.

**S9 Fig. Experimental cage set-up.** A) Schematic visualization of cage set-up in conventional Makrolon type III cages including running disc (mounted on igloo and pedestal), magnet, sensor, bike computer, nesting material, wooden enrichment , bedding and burrowing bottle (2 h per trial). B) Photo of running disc and igloo on pedestal, arrow pointing at magnet.

**S10 Fig. Nesting and grooming score.** A) Nest consolidation score using cotton rolls and B) grooming score (5).

**S1 Table. Parameters of modified Irwin test.** Scores in bold are considered physiologic.

### Supplemental data

#### Supplemental figure 1

A)

| Clinical Score | Activity | General condition | Behavior | Body posture | Body weight |
|---|---|---|---|---|---|
| 1 | Very active | Clear eyes; clean orifice | Alert, curios, species-specific | Normal, species-specific movements and posture | Weight gain; No change; Weight loss ≤ 5 % |
| 2 | Active | | | | Weight loss ≤ 10 % |
| 3A | Slightly reduced activity | Eyes partly closed | Alert, Hypo-/Hyperlocomotion, slightly reduced grooming | Normal, slightly curved dorsal line | Weight loss ≤ 15 % |
| 3B | Distinct reduced activity | Eyes partly closed | Frequent stops while movements, reduced food / water intake, reduced grooming | Slightly curved dorsal line | Weight loss ≤ 20 % |
| 4 | Slow | Eyes partly closed; dirty orifice | Reduced reaction to stimuli, absence of grooming behavior, functional loss of extremities | Curved dorsal line | Weight loss ≥ 20 % |
| 5 | Apathetic | Eyes closed; moist / clotted orifice | Self-isolation, no or negligible activity | Distinct curved dorsal line | Weight loss ≥ 20 % |
| 6 | Moribund | Eyes closed; flat breathing | No activity, no reaction to stimuli | Lateral position, seizures, animal is cold | Weight loss ≥ 20 % |

B)

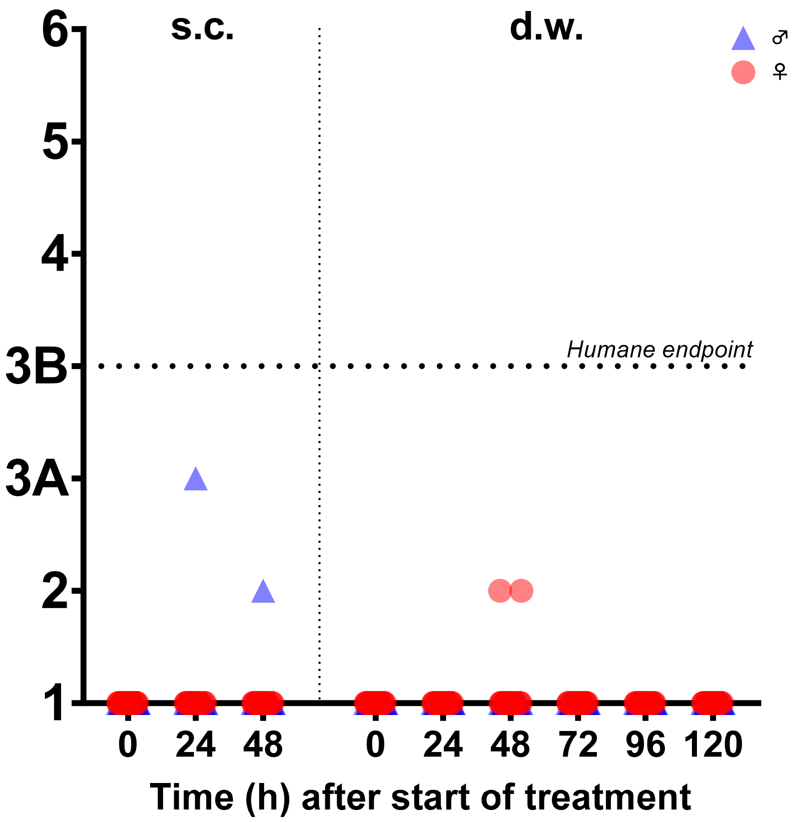

#### Supplemental figure 2

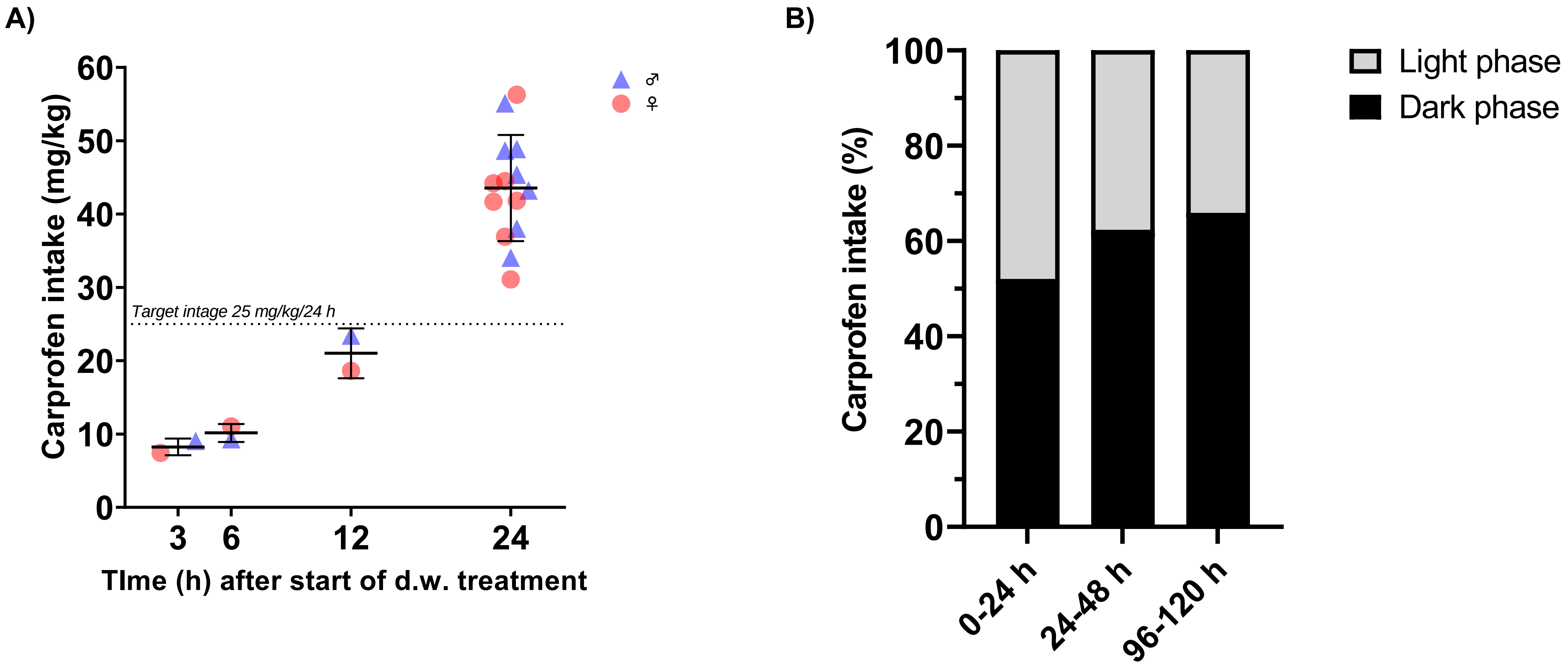

**Supplemental figure 3**

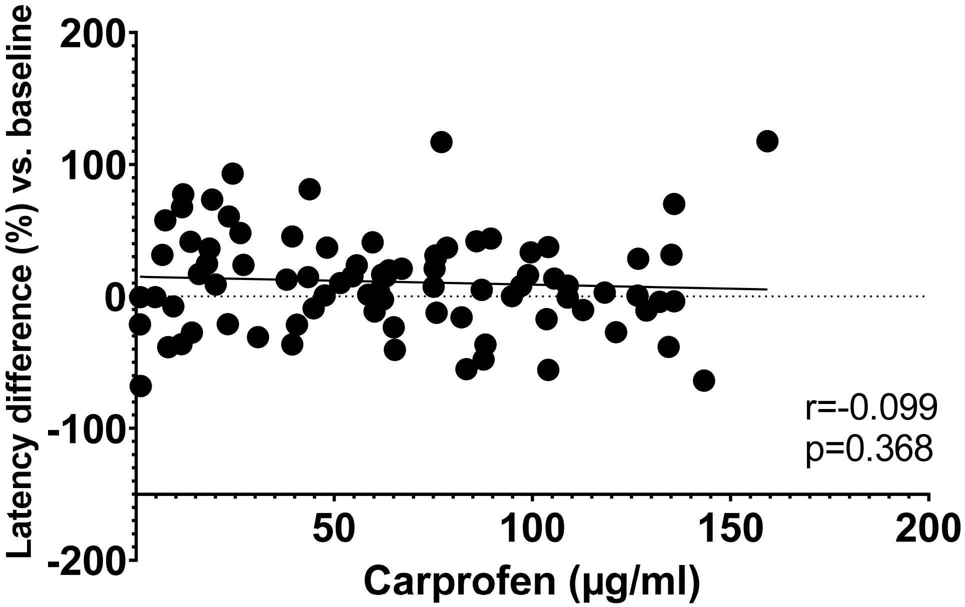

#### Supplemental figure 4

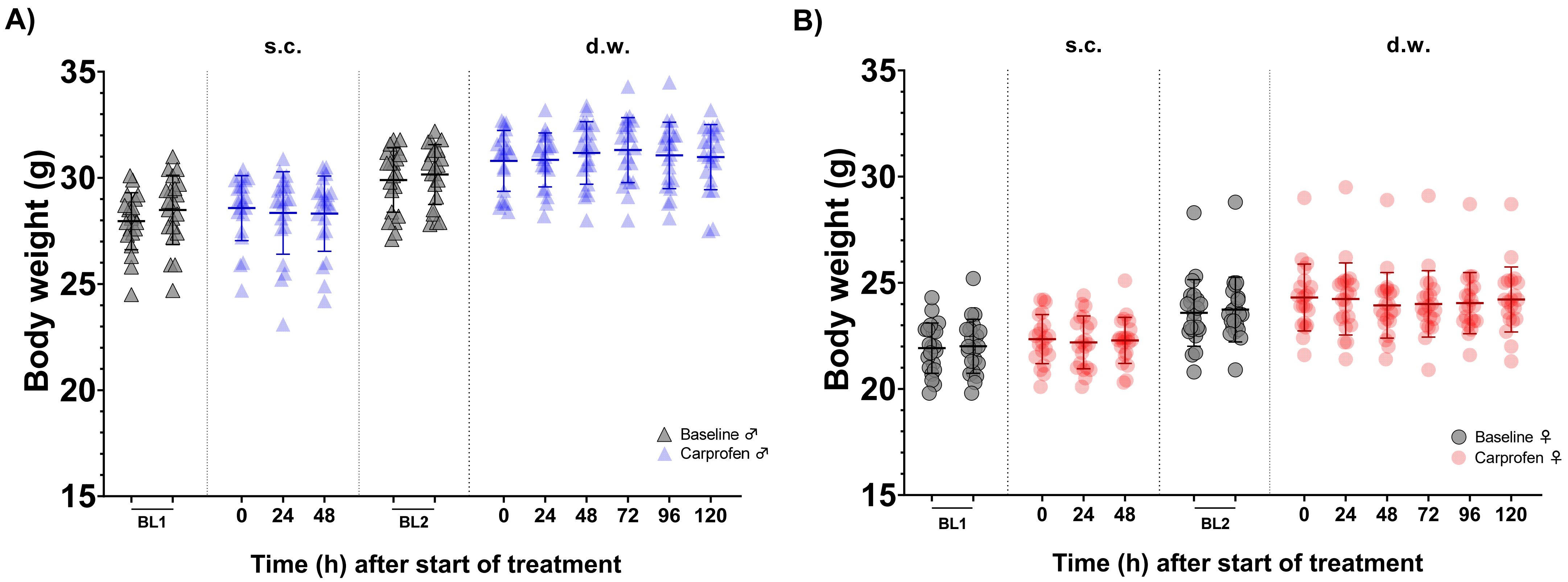

#### Supplemental figure 5

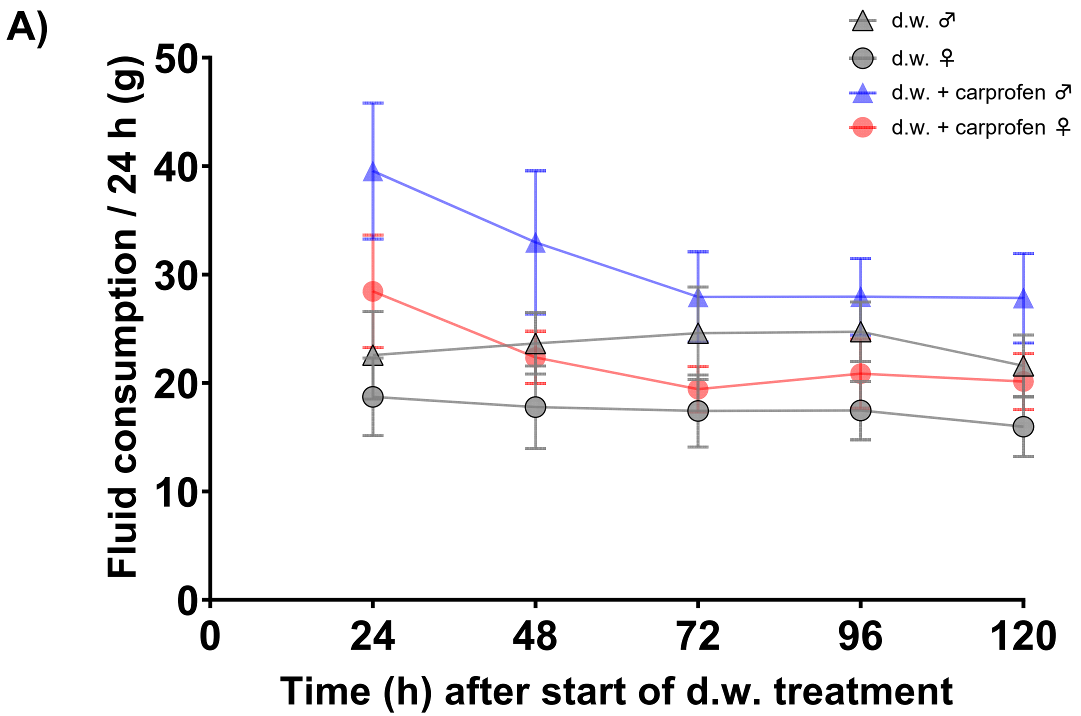


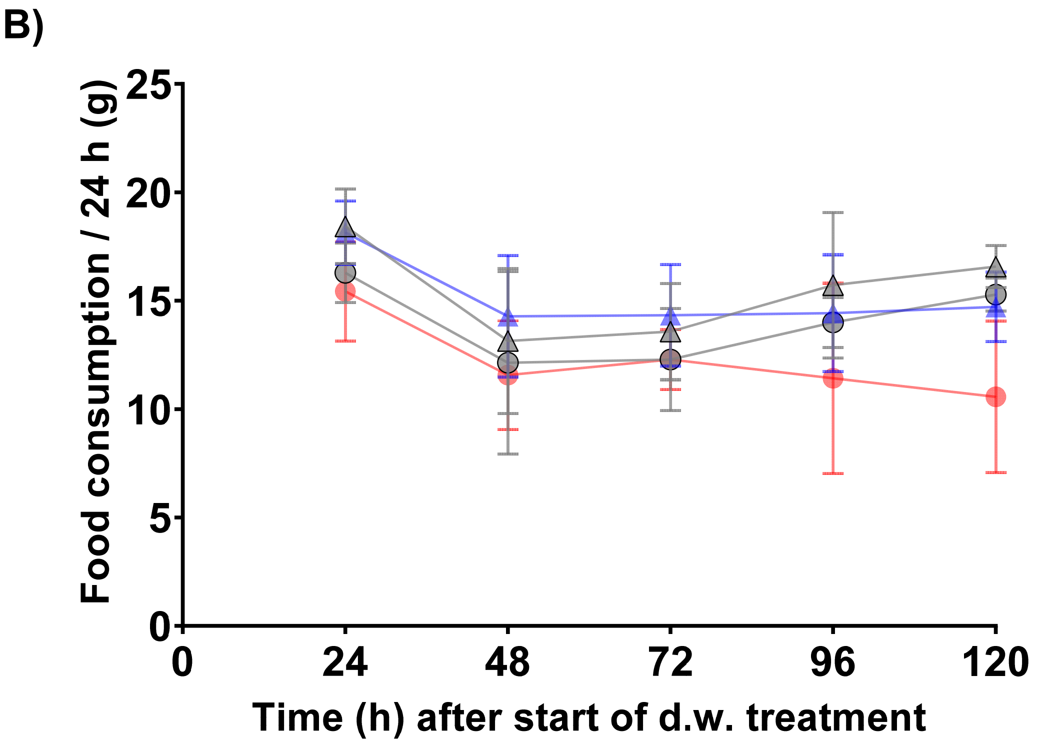

### Supplemental figure 6

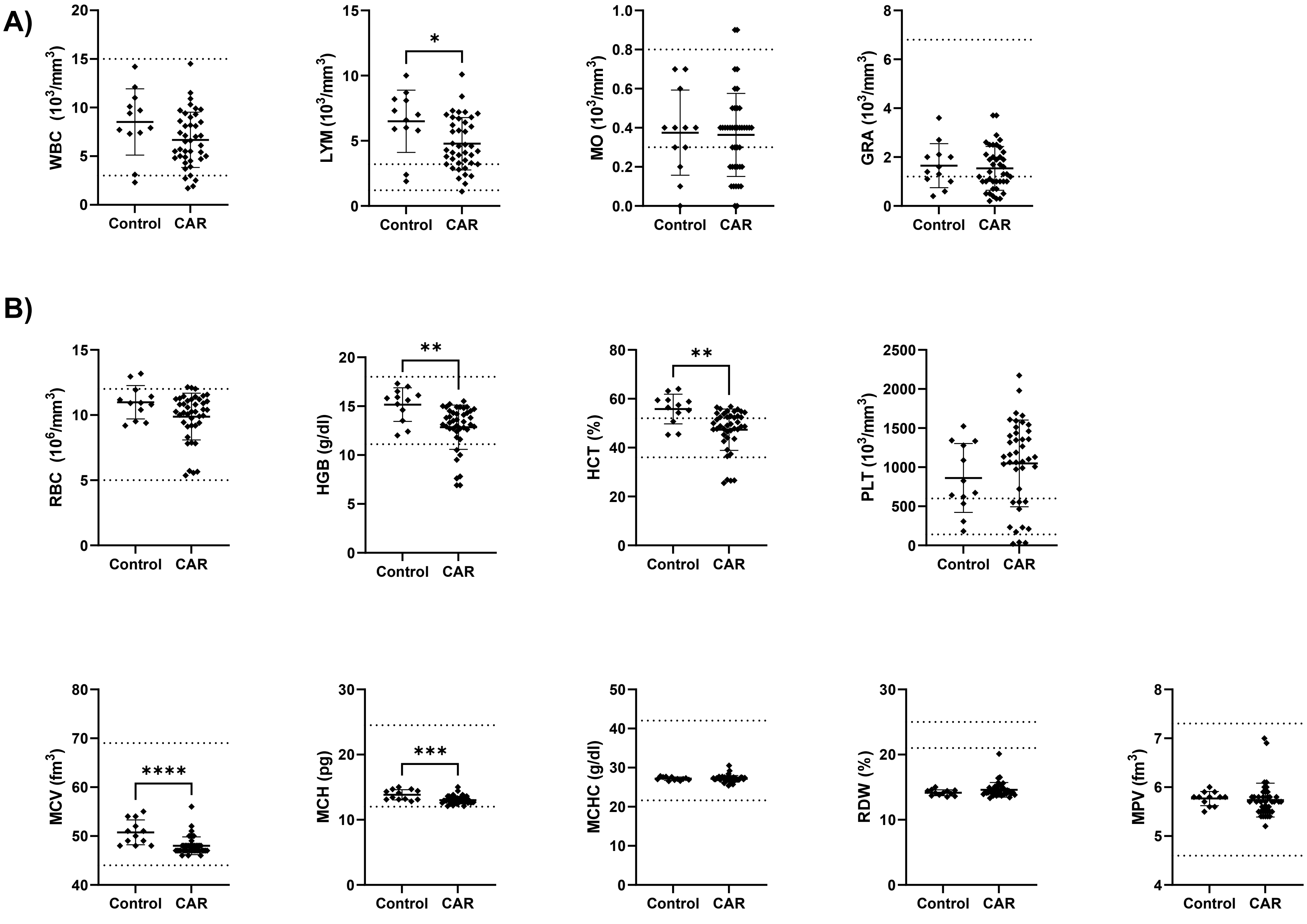

### Supplemental figure 7

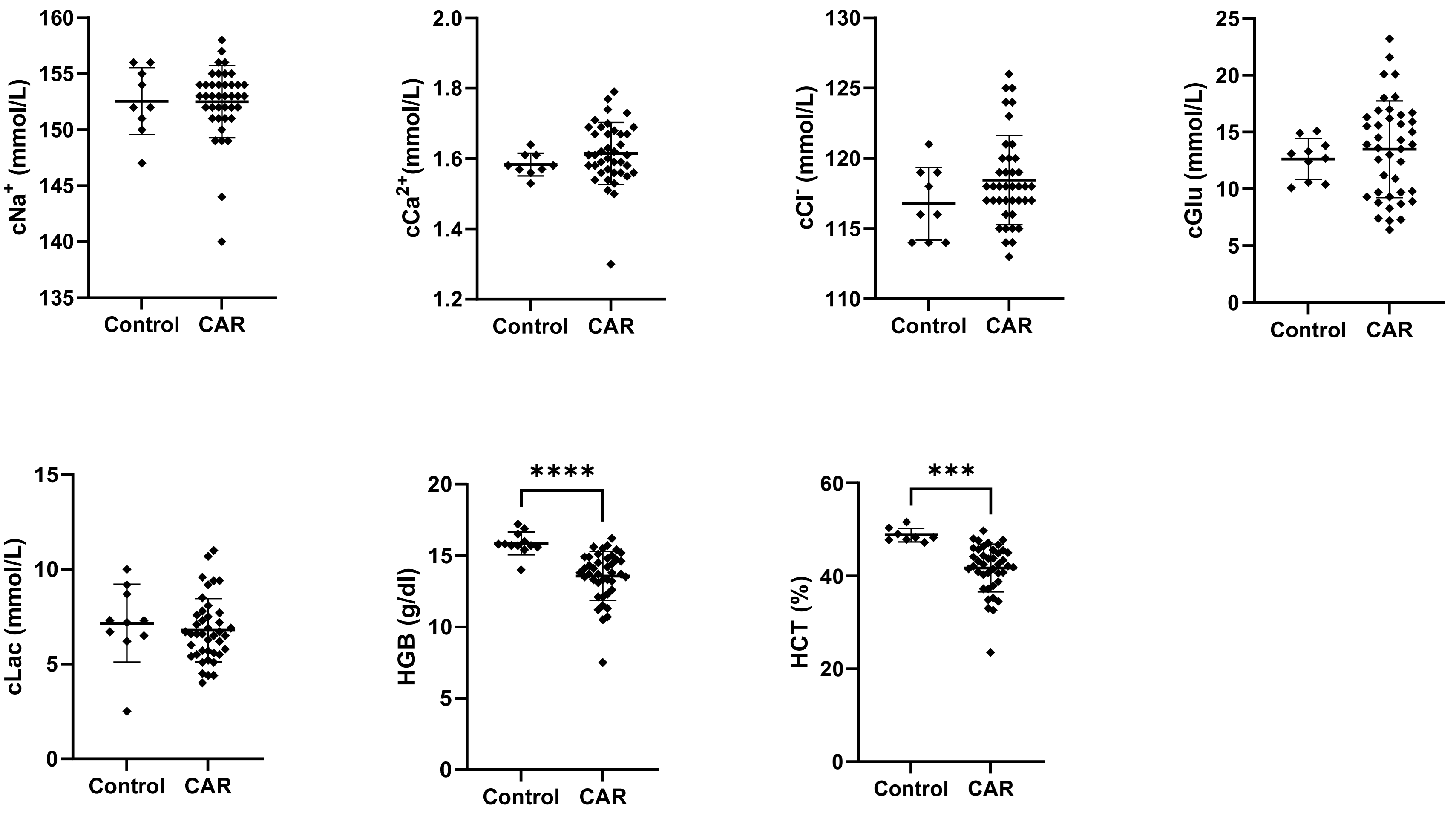

### Supplemental figure 8

A)

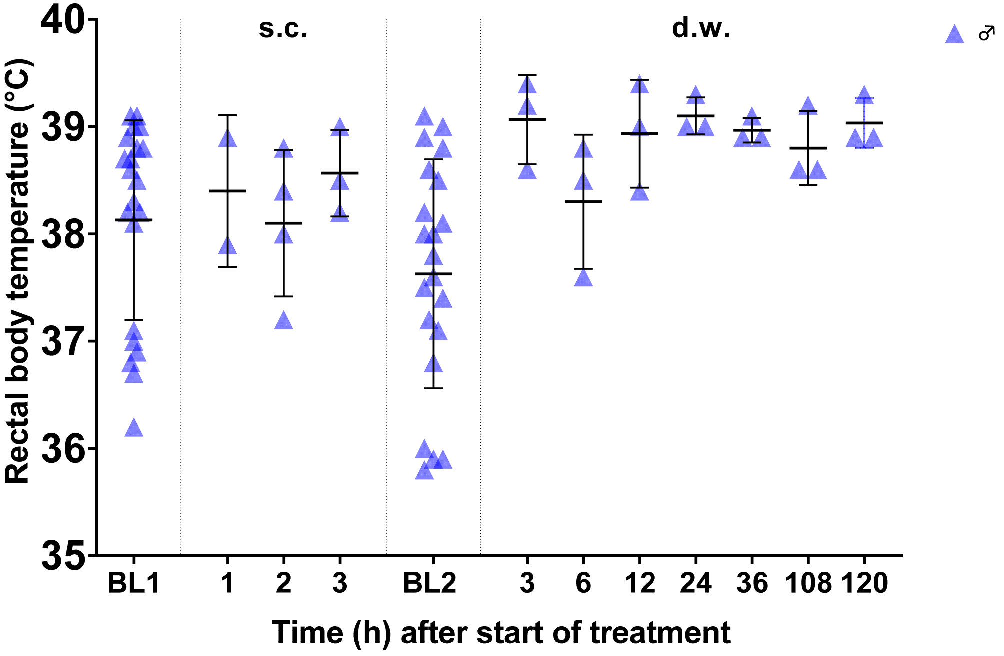


B)

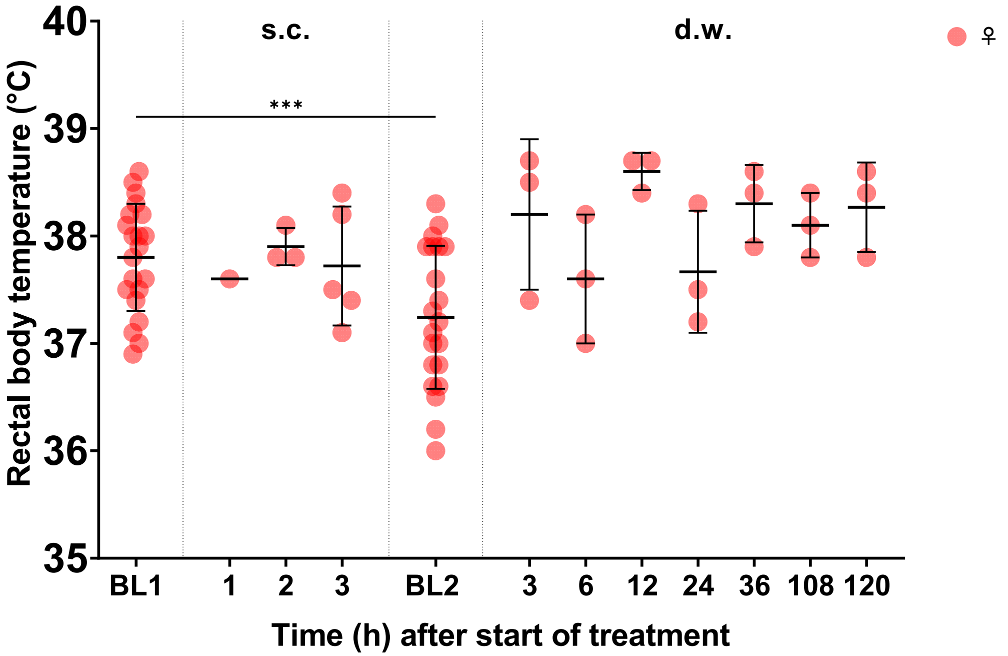

### Supplemental figure 9

A)

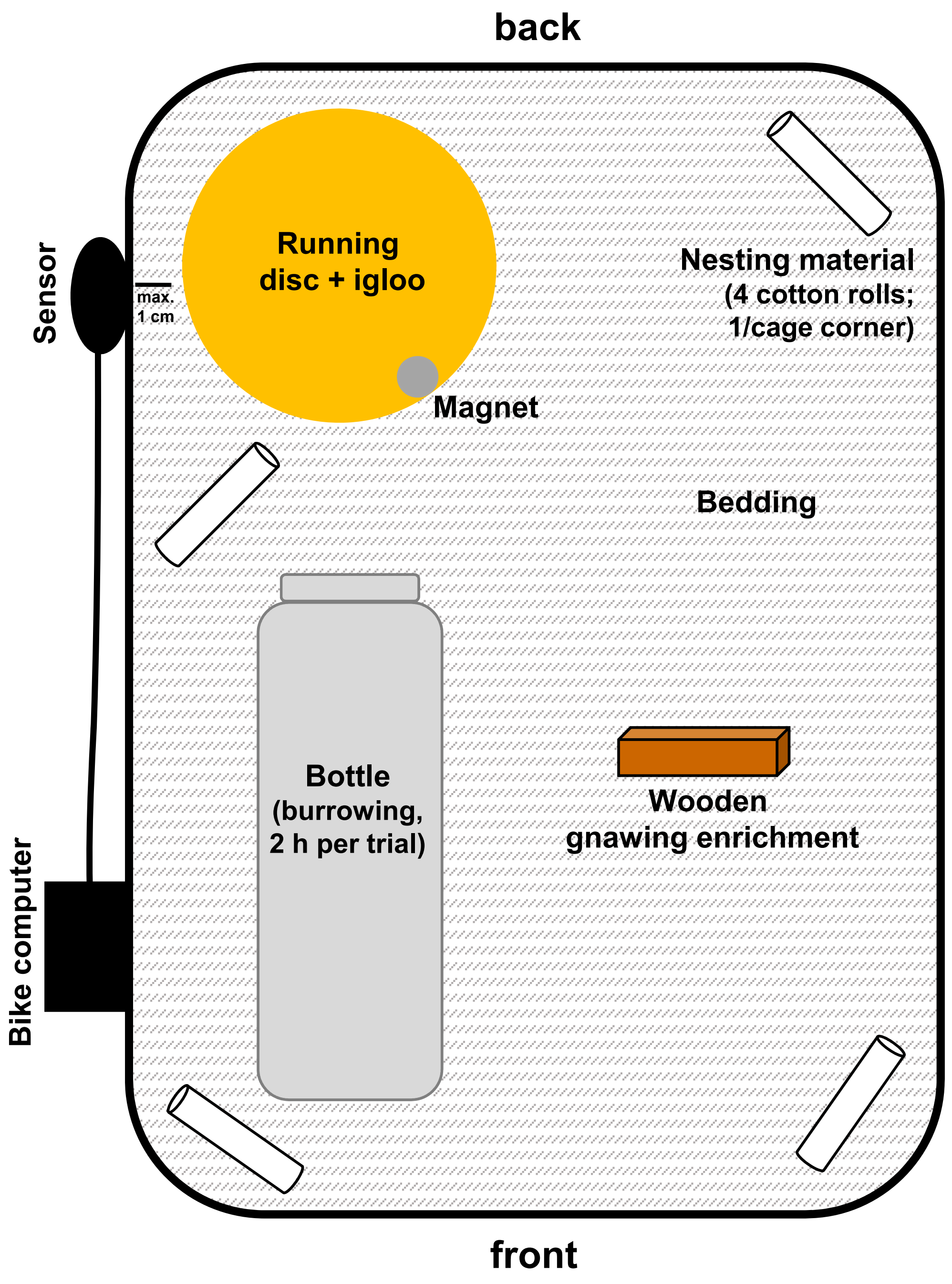


B)

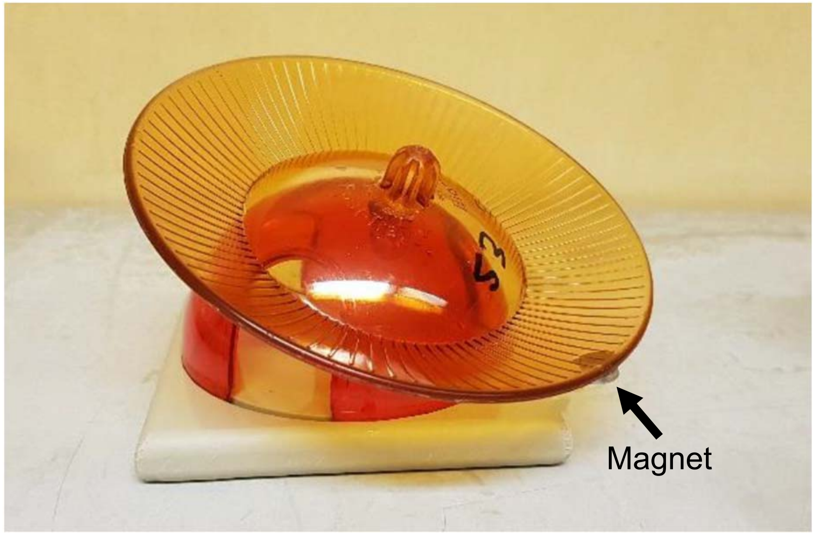

### Supplemental figure 10

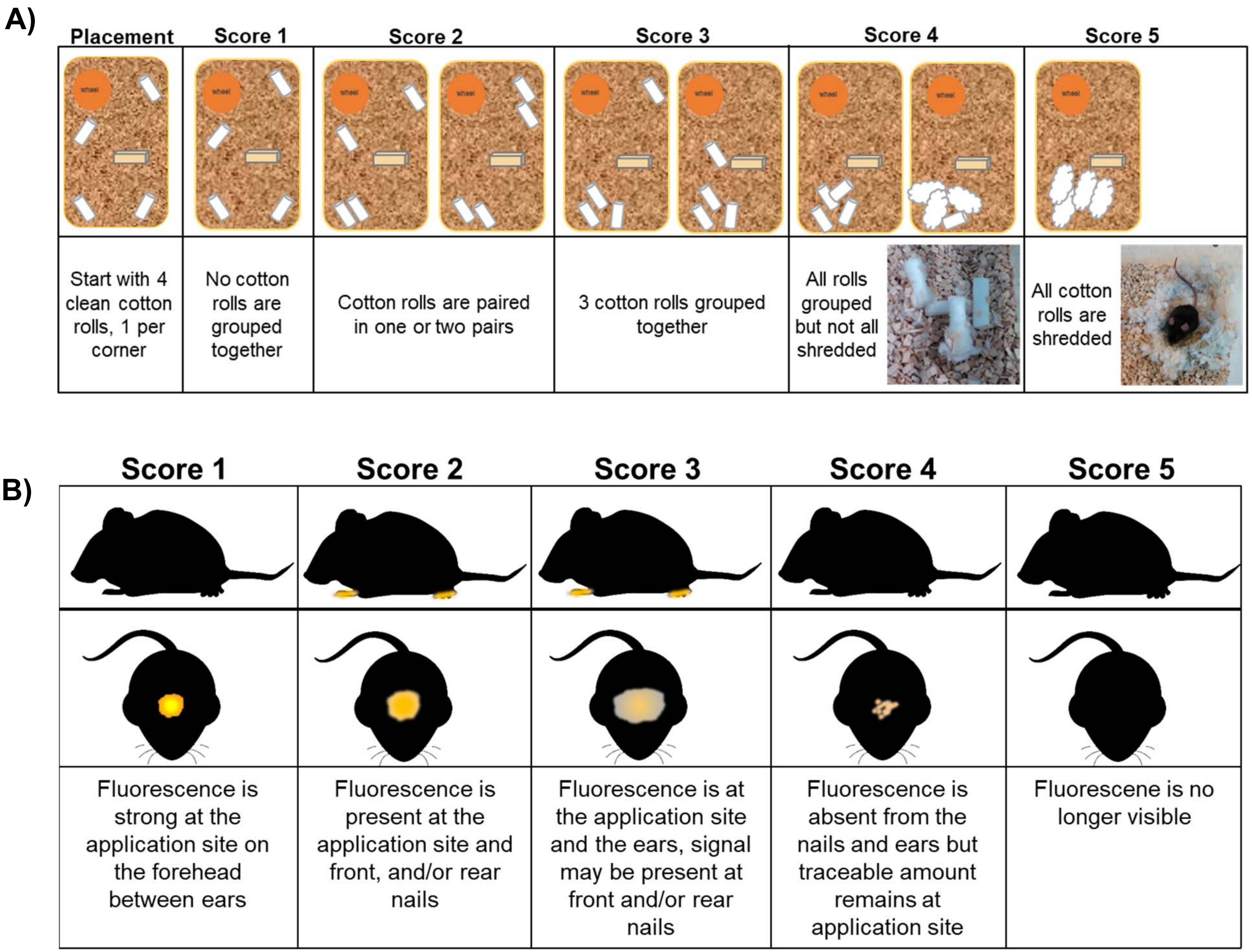
